# Mechanistic Basis of Sulfo Transfer in a PAPS-Independent ArylSulfate SulfoTransferases Revealed by Crystallography and QM/MM Simulations

**DOI:** 10.64898/2026.10.08.757636

**Authors:** Dilara Balci, Nina M. Nimbona, Haya Obeid, Aurélie Delavat, Lionel Cladière, Cyrille Grandjean, Mirjam Czjzek, Franck Daligault, Mehdi D. Davari

## Abstract

Enzymatic sulfation offers a green-chemistry route to the selective sulfation of phenolic and other hydroxylated compounds. Bacterial arylsulfate sulfotransferases (ASSTs) are particularly attractive biocatalysts because they use simple aryl sulfate donors, such as *p*-nitrophenyl sulfate (pNPS), instead of the complex and labile cofactor PAPS required by conventional sulfotransferases. However, their rational application and engineering remain limited because the molecular principles controlling substrate recognition, catalytically competent donor and acceptor positioning, covalent sulfo-enzyme formation, and sulfo transfer are not fully understood. In particular, it has remained unclear how conserved active-site histidines cooperate during catalysis and how active-site preorganization promotes efficient sulfo transfer. Here, we address these questions for the newly characterized ASST (ASTB) from *Desulfitobacterium hafniense* clade 6 (Dh_ASST_6) by combining X-ray crystallography, steady-state enzyme kinetics, and quantum mechanics/molecular mechanics (QM/MM) simulations. Crystal structures of Dh_ASST_6 in the apo state and donor-soaked state reveal a monomeric β-propeller enzyme with a histidine-centered active site and directly identify His350 as the transient sulfo carrier. Kinetic analysis shows efficient turnover and supports a ping-pong bi-bi mechanism, consistent with formation of a covalent sulfo-enzyme intermediate. Steered QM/MM simulations of the second catalytic step show that sulfuryl transfer from sulfated His350 to the phenolic acceptor proceeds through a proton-coupled, multistep pathway in which His283 assists acceptor activation by proton transfer. Active-site preorganization precedes the main bond-reorganization event and generates a near-attack geometry in which acceptor alignment, proton transfer, and stabilization of the sulfo group by a conserved hydrogen-bonding network jointly promote sulfo transfer. Free-energy reconstruction gives a low barrier of approximately 9.6 kcal mol⁻¹, in agreement with efficient enzymatic turnover. The structural, kinetic, and computational data support a complete catalytic cycle for Dh_ASST_6 based on a cooperative two-histidine mechanism. This model establishes catalytically competent acceptor positioning and preservation of the histidine-centered hydrogen-bonding network as key design principles for ASST engineering. These features provide concrete targets for expanding substrate scope and tuning donor or acceptor preference while maintaining efficient sulfuryl transfer.

## 1. Introduction

N- or O-sulfonation are widespread biochemical modifications that play important roles in metabolism, regulation and detoxification of diverse biomolecules and xenobiotics^1–3^. In general, these transformations involve transfer of a sulfo group (–SO₃⁻) to a substrate and can strongly affect its physiochemical and biological properties.^4–7^ In vivo, sulfation contributes to detoxification pathways^8^, hormone and steroid regulation^9^, and the processing of xenobiotics^8^ and natural products^10^. By converting hydrophobic compounds into more polar sulfate esters, it enables transport and excretion, while also serving regulatory functions in biosynthetic and signaling pathways^11, 12^. A wide variety of compounds can undergo N- or O-sulfonation^13^, including phenolics^14^, flavonoids^15, 16^, antibiotics^17^, steroids^18^, peptides^19^ and carbohydrates^20^, and in all these cases sulfation can influence solubility, stability and molecular recognition^21^.

From a synthetic and biocatalytic perspective, N- or O-sulfonations are highly attractive, yet challenging transformations^22, 23^. Conventional chemical approaches of N- or O-sulfonation often requires harsh reagents, protecting group strategies, and can lead to regioisomeric mixtures that are difficult to separate^22, 24^. In contrast, sulfotransferases (STs) catalyze sulfo-group transfer under mild, aqueous conditions and with high chemo-, regio- and often stereoselectivity^18, 25–29^. Their ability to selectively functionalize complex substrates makes them promising tools for late-stage modification of bioactive molecules^30^. Therefore, STs are promising tools for the preparation and late-stage functionalization of bioactive molecules.^31^

STs can be broadly divided into two classes according to the nature of their sulfo donor^25, 32–34^. Classical STs are PAPS-dependent enzymes that use 3’-phosphoadenosine-5’-phosphosulfate (PAPS) as an activated sulfo donor.^32^ Although highly effective in vivo, PAPS is structurally complex, relatively unstable and costly, limiting its applicability in biocatalysis^35^. In contrast, PAPS-independent STs, also known as arylsulfate sulfotransferases, (also termed aryl sulfotransferases) (ASSTs, EC 2.8.2.22), use simple aromatic sulfate esters such as *para*-nitrophenyl sulfate (*p*NPS) as sulfo donors^31, 33, 36, 37^.

ASSTs have therefore attracted considerable attention as biocatalysts for sulfo transfer reactions.^32^ This is mainly because they use simple aryl sulfate donors and because several ASSTs can accept a relatively broad range of substrates. More generally, enzymatic sulfation offers a green-chemistry route to the selective, mild, and in some cases monoselective sulfation of complex molecules. Although phenolic compounds are usually the preferred acceptors, some ASSTs can also act on more challenging non-phenolic acceptors, including linear alcohols and sugars, but generally with much lower activity than for phenolic compounds^18, 38–41^. This combination of donor accessibility and broad acceptor scope makes ASSTs especially appealing for the selective and mild sulfation of peptides, steroids, carbohydrates and other complex scaffolds, particularly in cases where PAPS-dependent STs are less suitable or chemical sulfonation lacks sufficient selectivity^18^.

Despite this potential, ASSTs remain far less characterized than PAPS-dependent STs.^10, 32, 42^ Only a limited number of ASSTs have been investigated in detail, although members of this enzyme family have been identified in a wide range of organisms, including pathogenic, commensal and environmental bacteria.^10, 32, 43^ This broad distribution already suggests that ASSTs are a diverse enzyme class with potentially distinct structural and functional subclasses^42^. Nevertheless, structural information is still limited^42, 44^. To date, only one ASST has been characterized by crystallography in detail, namely the enzyme from *Escherichia coli* (UniProt Q8FDI4, Ec_ASST_1), which revealed a distinct fold and identified key active-site residues involved in catalysis^33^. These studies established a general framework for PAPS-independent sulfo transfer but provide only a limited view of the structural and mechanistic diversity within the ASST family.^33^ This lack of structural knowledge also limits the rational engineering of ASSTs for biocatalytic applications, because the molecular features controlling activity, stability, and acceptor specificity remain poorly understood.

Available structural and kinetic data of ASSTs support a ping–pong bi–bi mechanism involving a covalent sulfo-enzyme intermediate^33, 44–46^. In this catalytic scheme, the sulfo donor first transfers its sulfo group to an active-site histidine residue, generating a transiently sulfonated enzyme^33, 44^. For the Ec_ASST_1, structural and kinetic studies identify His436 as the residue that undergoes transient sulfation during the first-half reaction, supporting strong experimental support for a histidine-centered mechanism^33, 44^. Although these findings clarify the identity of the sulfo-carrying residue, they do not explain how the second half-reaction proceeds. In particular, it remains unresolved whether additional conserved histidines and neighboring residues contribute directly to acceptor activation, proton transfer, or transition-state stabilization^32^. Understanding these roles is important because the second half-reaction determines how efficiently the sulfate group is transferred to different acceptor substrates. This is especially relevant for biocatalysis, where many non-phenolic acceptors, such as alcohols and sugars, are converted only with low activity despite their synthetic value. A clearer mechanistic picture would therefore not only deepen the fundamental understanding of how ASSTSs modulate sulfo-transfer reactivity but also provide a foundation for engineering enzymes with improved catalytic efficiency, broader acceptor scope, improved selectivity, and greater applicability in sustainable sulfation reactions.

In this context, ASTB from *Desulfitobacterium hafniense* clade 6 (Dh_ASST_6) provides an attractive system for mechanistic investigation, because *D. hafniense* arylsulfate sulfotransferases have already been developed and used repeatedly as practical biocatalysts for selective sulfation^18, 42, 47^. Enzymes from this organism are of particular interest because they can transfer sulfo groups not only to phenolic substrates but also to more challenging non-phenolic acceptors such as linear alcohols and sugars.^18, 41, 48^ This broader substrate scope is also reflected in successful applications of *D. hafniense* sulfotransferases to structurally diverse substrates, including catechols, quercetin derivatives, flavonolignans, luteolin, myricetin, ampelopsin, and related small molecules, showing their relevance from both mechanistic and biocatalytic perspectives.^15, 30, 49–51^ In addition, *D. hafniense* sulfotransferases have been used for regioselective sulfation of poorly soluble plant metabolites and in scalable chemoenzymatic sulfation workflows, further highlighting their synthetic utility.^14, 52^ Despite this biocatalytic relevance, molecular details of the catalytic mechanism in *D. hafniense* ASSTs remain insufficiently understood, including how donor and acceptor substrates are positioned in the active site and how conserved residues promote sulfo-group transfer.

Whereas the sulfo-enzyme intermediate has been firmly established, the mechanistic details of its turnover in the second half-reaction, considered the rate-limiting step, remain unknown. In particular, it is not known how the phenolic acceptor is positioned and activated in the active site, which residues assist proton transfer, how S-O bond formation is coupled to cleavage of the sulfo-enzyme intermediate, and which interactions stabilize the transition state.

Addressing these questions is important not only for understanding productive sulfo transfer in ASSTs, but also for guiding enzyme engineering campaign to improve catalytic performance, acceptor scope, and selectivity.

In this work, we report the X-ray crystallography of ASTB from *Desulfitobacterium hafniense.* The resulting structure provides a basis for examining the active-site architecture of ASTB and for comparing the arrangement of conserved catalytic residues with previously characterized ASSTs. We further performed steady-state kinetic analysis to characterize the catalytic activity of ASTB and provide experimental reference data for the computational part of the study. Molecular dynamics simulations were then used to examine how the sulfo-enzyme intermediate and phenol are arranged in the active site before the second catalytic step. Building on these structures, we applied QM/MM free energy simulations to investigate how the sulfo group is transferred from the sulfo-histidine intermediate to phenol. This combined approach allowed us to address how phenol is activated, how proton transfer is coupled to sulfo transfer, and how the sulfo-histidine bond is cleaved during turnover. The computed free-energy profile is consistent with kinetic data, supporting the assignment of the second half-reaction as the rate-limiting step. These simulations further suggest that phenol is first activated by proton transfer to a conserved histidine, before the resulting activated acceptor attacks the sulfo group. These findings allow us to propose a complete catalytic reaction cycle for ASTB and provide a molecular explanation for how acceptor activation is coupled to productive sulfo transfer in ASSTs.

## 2. Methods

### 2.1. Protein Purification

Dh_ASST_6 was produced in 300 mL of auto-induction ZYP 5052 medium supplemented with 100 µg·mL^-1^ ampicillin. Cultures were grown at 20°C with 150 rpm for 72 hours. Cells were harvested by centrifugation at 5000×g, 4°C for 45 minutes. The supernatants were discarded and pellets were resuspended in 20 mM HEPES (pH 7.5), 300 mM NaCl and disrupted by sonication. The cell lysates were centrifuged at 20,000×g, 4°C and for 45 minutes. The soluble fraction was then purified by affinity chromatography on Äkta system using a 5 mL Histrap FF (Cytiva) and equilibrated with 20 mM (pH 7.5) HEPES buffer containing 300 mM NaCl and 20 mM imidazole. This step was followed by size exclusion chromatography using a HiLoad 16/600 Superdex 200pg column (Cytiva), equilibrated with 20 mM MES buffer (pH 6.5) containing 200 mM NaCl. The purified protein was concentrated using a 30 kDa molecular-weight cutoff to a final concentration of 34 mg·mL^-1^.

### 2.2. Protein Crystallization and Structure Determination

Crystallization experiments were performed by sitting-drop vapor diffusion using a Crystal Gryphon nanodrop robot (Art Robbins Instruments) and commercial crystallization screens including PACT Premier Eco Screen, PEGsuite, PEGsuite II, and JCSG-plus Eco Screen. Initial crystallization conditions for DH_ASST_6 were then manually optimized in hanging drops, leading to best conditions by mixing 2 µL of enzyme solution (17 mg·mL⁻¹) with 2 µL of reservoir solution containing MES (pH 6.5): 0.18 M and PEG 10,000: 17% at 16 °C. For the Dh_ASST_6_*p*NPS complex, native crystals were soaked 5 min directly in the crystallization drop supplemented with *p*-nitrophenyl sulfate (*p*NPS) as powder. Crystals were flash-frozen in liquid nitrogen using 20% (v/v) glycerol as cryoprotectant. Diffraction data for native Dh_ASST_6 alone and in complex with *p*NPS were collected at the PROXIMA1 beamline, synchrotron SOLEIL (Paris, France). Data processing was carried out with XDS^53^ and data were scaled with Aimless (CCP4 package)^54^. The apo-structure was solved by molecular replacement using Phaser^55^ from the CCP4 and an AlphaFold2-predicted^56^ model of Dh_ASST_6 (UniProt: B8FZF6) as search template, while the substrate complex was solved using Dh_ASST_6_apo as starting point. Refinement was performed using REFMAC5^57^, with iterative model building in COOT^58^. Water molecules were automatically added with WARP (integrated in REFMAC5) and manually confirmed or deleted. Structure validation was performed with MolProbity^59^. Data collection and refinement statistics are given in **Table S1,** and final coordinates were deposited in the Protein Data Bank under accession codes 30FK and 30FV.

### 2.3. Enzymatic Assays and Kinetic Analysis

Optimal assay conditions for Dh_ASST_6 activity were established at pH 8.5 and 39 °C and were used for all subsequent enzymatic assays (data not shown). Reactions were carried out in 96-well microplates in a final volume of 200 µL containing 50 mM AMPSO buffer (pH 8.5) and 13.8 nM Dh_ASST_6. Enzymatic activity was monitored continuously at 405 nm using a Tecan Infinite M1000 microplate reader at 39 °C for 30 min. Initial velocities were determined from the slope of the initial linear phase of the absorbance-time traces.

Steady-state kinetic parameters were determined using *p*NPS and phenol as substrates. For the kinetic analysis, *p*NPS concentrations of 0.14, 0.5, 1, 2, 5, 8, 12, and 15 mM were used, while phenol concentrations of 0.25, 0.5, 3, 8, 10, 20, 30, 40, 80, and 100 mM were tested. Kinetic data were analyzed using GraphPad Prism 8.0.1. Initial-rate data were fitted to a bi-substrate kinetic model according to the following equation:

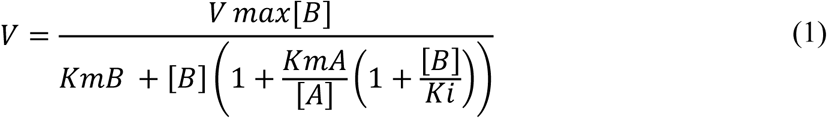

Double-reciprocal Lineweaver–Burk plots were generated by plotting 1/V_i_ against 1/[*p*NPS]. Secondary replots of slopes and y-intercepts were used to determine the kinetic constants *K*_m,*p*NS_, *K*_m,phenol_, *K*_i,phenol_, and V_max_.

### 2.4. Computational Methods

#### 2.4.1. Protein Loop Modelling

The resolved crystal structure of Dh_ASST_6 (PDB 30FK) lacked electron density for amino acid residues 576–590. This missing segment was modeled and conformationally sampled using YASARA v.23.9.23^60^ by the loop-building routine, in which candidate loop conformations compatible with the flanking anchor residues are generated from knowledge-based structural templates. All generated loop models were subsequently energy-minimized using the steepest-descent algorithm. During minimization, the modeled loop residues together with one adjacent residue on each side were allowed to move, while the remainder of the protein was kept fixed. To account for structural variability among the generated models, including differences in loop orientation and conformation, the minimized structures were clustered with a custom Python script using hierarchical clustering based on loop RMSD. Representative structures from each cluster were then selected as initial structure for short molecular dynamics simulations. The unresolved N-terminal residue, Met1, was added manually in PyMOL v3.0^61^.

#### 2.4.2. Protein Structural and Bioinformatic Analysis

The crystal structure of Dh_ASST_6 (UniProt: B8FZF6, PDB 30FK) was used as the starting model for structural analysis. For all subsequent calculations, only chain A was kept. Conserved residues were identified using the UET web server^62^, utilizing as input the crystal structure (PDB code 30FK) in which missing loop regions had been modeled. In addition, sequence homologs were identified by BLAST^63^ searches using the Dh_ASST_6 sequence as query. The resulting multiple-sequence alignment was visualized and analyzed with the NCBI Multiple Sequence Alignment Viewer v1.26.0^64^ to inspect evolutionary conserved sequence regions and active-site residues, with particular attention to conserved histidines located in the catalytic center. Structural analysis of the modelled enzyme was carried out using CAVER v3.0^65^ to identify tunnels connecting the protein surface with the active-site region and KVFinder^66^ to characterize the geometry and extent of the catalytic cavity.

#### 2.4.3. Structure-Guided Substrate Placement of Phenol

Phenol was manually placed into the active site of Dh_ASST_6 in PyMOL^61^ using a structure-guided model building approach. The starting model represented the sulfo-enzyme intermediate, in which the catalytic histidine was covalently linked to the sulfo group, corresponding to the state before the second half-reaction. Chain A of Ec_ASST_1^33^ (PDB 3ETT; 2.10 Å resolution) served as the reference structure because it contains a bound aryl ligand (4-nitrophenol) in a comparable catalytic pocket. Following structural superposition of Dh_ASST_6 with Ec_ASST_1^33^, phenol was positioned in the active site defined by His220, His283, Arg301, and His350, using the ligand coordinates in the reference structure as a guide. The six shared aromatic carbon atoms were then used for pairwise ligand fitting; the nitro substituent of 4-nitrophenol was excluded. The resulting pose positioned the phenolic hydroxyl group near His283 and the sulfo group on His350. The model was inspected for steric compatibility and used as the starting sulfo-enzyme–acceptor complex for MD. This procedure constitutes alignment-guided model construction; no automated pose search, docking score, or binding-energy ranking was performed (Section S3.5).

#### 2.4.4. Constant pH molecular dynamics Simulation

Constant pH molecular dynamics (CpHMD) in implicit solvent^67^ was performed with Amber22^68^ and ff10 force field to examine His220, His283, and His350 in the substrate-free model. These three residues were initially assigned the doubly protonated HIP state and were the only titratable sites. The system underwent restrained heavy-atom minimization, restrained whole-protein relaxation, and unrestrained minimization, followed by heating from 100 to 200 K and from 200 to 300 K with a 0.5 fs time step under Langevin dynamics. Production simulations covered the reported pH range 0–7 and comprised 1,000,000 steps of 2 fs (2 ns) at 300 K SHAKE^69^ constrained bonds involving hydrogen, and Monte Carlo moves sampled protonation states. The populations discussed here are those reported at pH 6.5 in Section S3.4 and **Table S7**. The assignment used for His283 in the second half-reaction model was subsequently changed to neutral HID for the mechanistic reasons discussed in Section 3.4.

#### 2.4.5. Molecular Dynamics Simulations

Except for His220, His283, and sulfo-His350, which were assigned as described above, protonation states of titratable protein residues were determined with the H++ server^70^ at pH 6.5. All Molecular Dynamics (MD) simulations were performed using Amber22^68^ with the ff19SB^71^ force field parameters for the enzyme. The acceptor and sulfated His350 were defined using AMBER/GAFF^72^ atom types. Missing point charges were calculated through QM optimization (B3LYP/6-31+G*)^73–77^ in Gaussian16^78^ and were parameterized using a RESP^79^ model (RHF/6-31G**)^75, 76, 80^ on the optimized geometry with Antechamber^72^. The systems were solvated under periodic boundary conditions in a cubic simulation box with OPC^81^ water molecules and neutralized with Na+ ions. The system underwent a three-step energy minimization process: initially relaxing protons, followed by the solvent, and finally the entire system. Next, it was heated in two steps with a linear increase of the temperature and a harmonic force on the backbone residues of the enzyme. The first heating step was conducted from 0 to 200 K and the second from 200 to 300 K in 1 ns using the Langevin thermostat in a NVT ensemble. The equilibration procedure consists of six 20 ps steps with gradually removing the restraints. The first four steps were NVT equilibration, while the remaining two steps were NPT ensembles, with no restraints on the backbone atoms of the enzyme. Six simulations were conducted with a time step of 2 fs for 200 ns with periodic boundary conditions and a cutoff of 10 Å was defined for the non-bonded interactions. All water molecules were processed using the SHAKE^69^ algorithm. Trajectories were analyzed with cpptraj^82^ and the python package MDAnalysis^83^.

#### 2.4.6. Steered QM/MM MD Simulations

Steered QM/MM MD simulations of the second half-reaction were performed with sander in Amber22^68^. Phenol was neutral, and His283 was assigned the neutral HID tautomer (Nδ1–H, with Nε2 available for proton acceptance). The expanded QM region comprised phenol, the sulfo group, and side chains of His220, His283, Arg301, His350, Tyr194, Asn285, Tyr416, Ile417, and Tyr487, with a total QM-region charge of +1. Amino-acid residues were truncated at Cα–Cβ bonds and capped with hydrogen link atoms^84^, phenol was retained intact. The QM subsystem was described at the DFTB3^85^ level with the 3OB parameter set, which has been developed and parameterized for organic and biological systems^86^, including sulfur-containing compounds^87^, and has been used in QM/MM studies of enzyme-catalyzed reactions^88, 89^. Additionally, a QM/MM cutoff of 10 Å was applied. To drive the reaction, two distance-based collective variables (CV) were used. CV1 was the phenolic H···Nε2(His283) distance and CV2 the phenolic O···S(sulfo-His350) distance. Both were shortened simultaneously using linearly moving harmonic restraints with a force constant of 600 kcal mol⁻¹ Å⁻². Each trajectory lasted 2 ps at 300 K, with a 0.2 fs time step, Langevin dynamics, and periodic boundary conditions. The 314 starting frames were selected from classical MD using O···S < 5.5 Å and H···Nε2 < 2.2 Å. QM-region and CV tests are described in Section S4 and **Table S8**.

The reported work-derived profile was reconstructed using the Jarzynski exponential-work estimator^90^(eq 2). At a given steering time, the average is over trajectories, using the work accumulated up to that time:

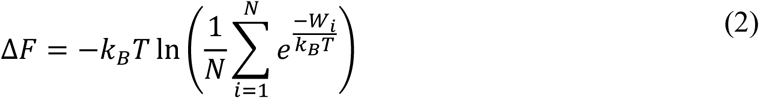

Here, Wᵢ is the accumulated work in trajectory i up to the selected steering time, N is the number of trajectories, kB is the Boltzmann constant, and T is temperature. Thermal energy and work must be expressed in consistent units; for molar work, the corresponding thermal factor is RT. Correlated frames from one trajectory do not constitute additional independent work realizations. The value at the end of the protocol uses the final accumulated work, whereas a time-resolved profile requires the accumulated work at every reported time. Equation 2 relates equilibrium free energies of the Hamiltonians defined by the steering protocol when its ensemble assumptions are satisfied; reconstructing an unbiased molecular free-energy landscape additionally requires treatment of the applied bias. We therefore refer to the plotted quantity as a work-derived profile and discuss its interpretation and sampling limitations in Section 3.5.3.

## 3. Results and Discussion

### 3.1. Crystal Structure of *Desulfitobacterium hafniense* clade 6 (Dh_ASST_6) Reveal a Preorganized Active Site and Capture the Covalent Sulfo-Enzyme Intermediate

The structures of native Dh_ASST_6 and *p*NPS-soaked Dh_ASST_6 establish a conserved catalytic scaffold and capture two distinct sulfo-binding states. The structures were determined at 1.90 and 2.79 Å resolution, respectively, in space group P2₁2₁2₁, with two molecules in each asymmetric unit (**Table S1**). The limited intermolecular contact area of 594 Å², together with a single size-exclusion peak consistent with a monomer, supports interpretation of the crystallographic pair as a packing contact. Each monomer contains an N-terminal domain (residues 1–118), a central six-bladed β-propeller catalytic domain (119–513), and a C-terminal domain (514–626). The extra domains have β-sandwich-like architectures (**Figure 1a**). Residues 576–590 near the pocket entrance were unresolved in both structures. The larger segment 575–591 used in subsequent loop displays and RMSD analysis includes the two flanking anchor residues.

**Figure 1.**
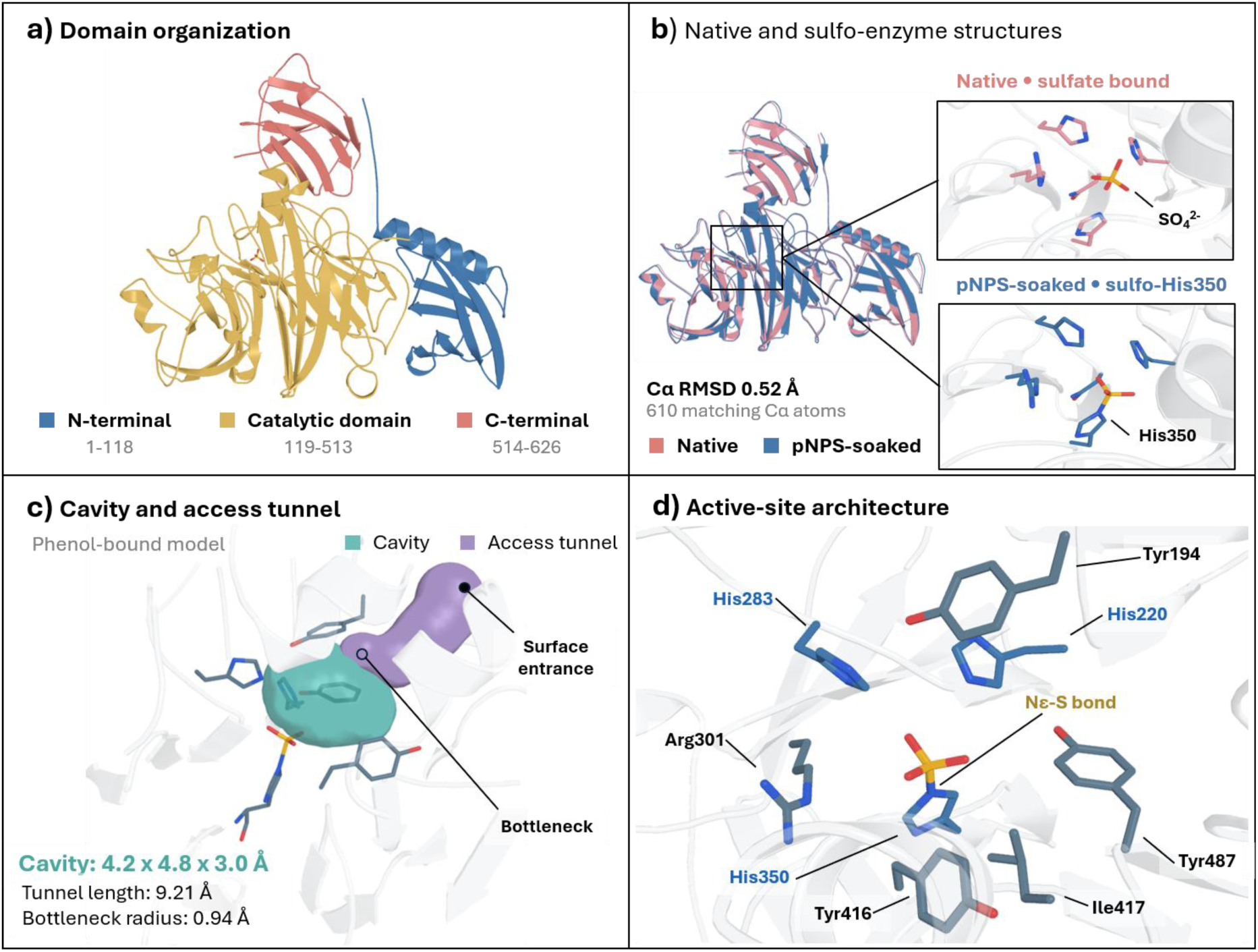
Structural organization and active-site architecture of Dh_ASST_6. **a)** O Monomer with the N-terminal domain in blue, the catalytic β-propeller in gold, and the C-terminal domain in coral. **b)** Superposition of Dh_ASST_6 (pink) and the Dh_ASST_6_*p*NPS (blue), highlighting the conserved overall fold and the active-site region in which the transient sulfated histidine intermediate is trapped at His350**. c)** Representation of the active-site cavity and the preferred access tunnel leading to the catalytic pocket. **d)** Close-up view of the active site with key catalytic residues labeled, including Tyr194, His220, His283, Arg301, His350, Tyr416, Ile417, and Tyr487.

Superposition of the native and donor-soaked structures gives an RMSD of 0.52 Å over 610 Cα atoms (**Figure 1b**). Sulfo-enzyme formation is therefore compatible with preservation of the overall scaffold. This structural similarity supports a prearranged catalytic core but does not establish that local side-chain motions, protonation changes, or substrate reorientation are energetically negligible. His220, His283, and His350 occupy the base of the β-propeller cavity, surrounded by Arg301, Asn285, Tyr194, Tyr416, Ile417, and Tyr487 (**Figure 1d**). This arrangement provides a framework for separating the chemistry of the conserved center from the less conserved features that control substrate access and orientation.

The native structure contains density assigned to a sulfate ion and is therefore more precisely described as native/sulfate-bound than ligand-free. In the *p*NPS-soaked structure, the sulfo group is covalently attached to His350 Nε2. This observation directly identifies His350 as the sulfo carrier, rather than assigning this function only by sequence similarity. The surrounding polar contacts are largely retained between the two structures, proposing that the pocket accommodates both noncovalent sulfate binding and the covalent intermediate without a large rearrangement of the backbone. The structural evidence is strongest for the identity of the carrier. The roles of the other histidines in proton transfer require additional mechanistic evidence.

Conservation analysis supports the importance of this catalytic environment while distinguishing direct contacts from pocket-shaping residues (Section S3.1; **Figures S2** and **S3**; **Table S**3). Tyr194, His220, His283, Asn285, Arg301, and His350 are conserved in the 100-sequence alignment. Tyr416 and Ile417 are conserved in all sequences covering those positions, whereas Tyr487 permits a conservative Tyr-to-Phe substitution. In the native structure, the histidines, Asn285, and the Ile417 backbone make short sulfate contacts of approximately 2.6–3.1 Å. Several conserved aromatic residues are farther from sulfate and instead define the surrounding pocket.

The cavity dimensions reported for the displayed model, approximately 4.2 × 4.8 × 3.0 Å, are consistent with a compact substrate-binding region (**Figure 1c**). The phenol-bound short-route analysis shown in that panel reports a length of 9.21 Å and a bottleneck radius of 0.94 Å. A separate CAVER analysis of the loop-modelled structure identifies 29 pathways in eight clusters, whose representatives are 23.90–62.85 Å long with bottleneck radii of 0.704–0.746 Å (Section S3.3; **Tables S5** and **S6**; **Figure S5**). These analyses describe different sets of geometric paths and should not be treated as interchangeable measurements of one tunnel.

Comparison with Ec_ASST_1 places these results in the context of the established ASST mechanism. Even though approximately 25% sequence identity, the catalytic histidines occupy corresponding positions, consistent with a conserved sulfo-transfer core (Section S2.1; **Figure S1**). The wild-type Ec_ASST_1 sulfo-enzyme structure contains the donor product 4-nitrophenol (PDB 3ETT), whereas no corresponding product density was observed in donor-soaked Dh_ASST_6. Differences near Tyr194, Tyr416, and Ile417 could alter aromatic packing and ligand orientation. However, absence of product density cannot by itself distinguish weaker binding, faster release, incomplete occupancy, or differences in crystallization conditions. The comparison supports variation in the substrate-binding environment without establishing that Dh_ASST_6 is intrinsically less rigid or has a broader substrate scope than Ec_ASST_1.

### 3.2. Enzyme Kinetics Analysis Reveals Ping-Pong Catalysis through a Transient Sulfo-Enzyme Intermediate

ASSTs are bireactant enzymes described as bi bi systems (bisubstrate-biproduct) with two substrates (a sulfo group donor, *p*NPS, and an acceptor, phenol) and two products (the desulfonated donor, *p*NP, and the sulfonated acceptor, phenol sulfate).^91^ This type of system can follow different mechanisms such as the sequential mechanism (ordered or random) or the ping-pong mechanism. These different mechanisms can be distinguished using double reciprocal Lineweaver-Burk plots, which show intersecting or parallel lines in sequential or ping pong mechanisms respectively, when the concentration of one substrate is varied whilst keeping the other constant.^92^ Steady-state kinetic experiments were performed to determine the type of mechanism of Dh_ASST_6 (ASTB). A series of experiments was conducted by varying the concentrations of *p*NPS and phenol under conditions ranging from undersaturation to saturation. **Figure 2a** shows the plot of the initial rate of the reaction catalyzed by Dh_ASST_6 and highlights substrate inhibition at high concentrations of phenol, which competes with *p*NPS at the active site. This inhibition was overcome at a higher concentration of *p*NPS (1 mM). Plots of the reciprocal Lineweaver Burke representations of 1/Vi versus 1/[*p*NPS] yielded a set of parallel lines at low phenol concentrations and another set of intersecting lines at higher phenol concentrations (**Figure 2b**). All these data are characteristic of ping-pong kinetics with competitive substrate inhibition by the acceptor.^1^ True kinetic constants were extracted from secondary replots of slope of 1/[*p*NPS] and 1/V_max,app_ versus [phenol] (**Figure 2c**): *K*_M,*p*NPS_ = 2.25 x 10^-4^ M, *K*_M,phenol_ = 4.23 x 10^-4^ M, *K*_I,phenol_ = 4.515 x 10^-2^ M, and *k*_cat_ = 14.5 s^-1^. These results are consistent with the formation of a transient sulfo-enzyme intermediate.

**Figure 2.**
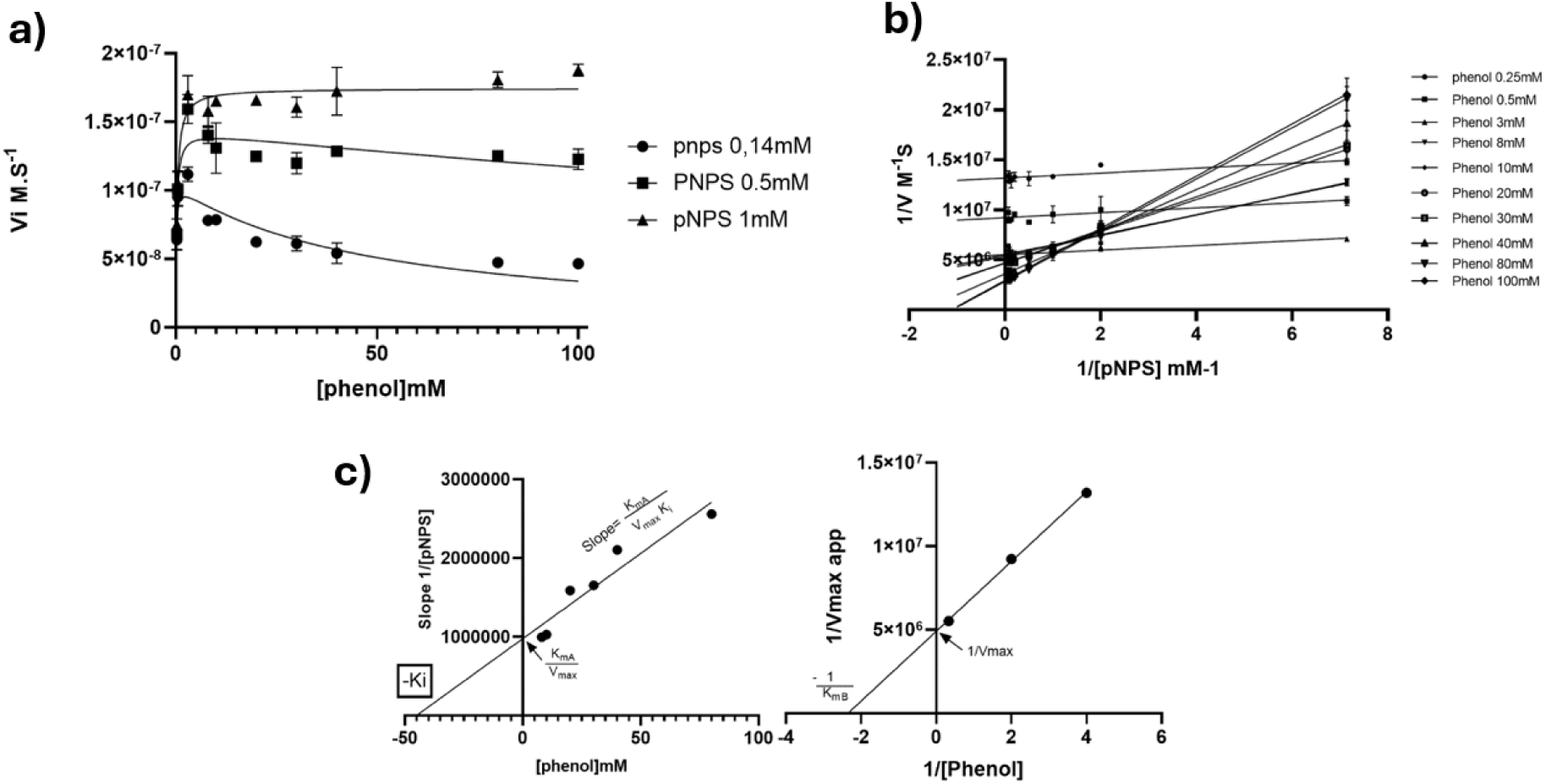
Kinetic analysis of ASTB catalyzed sulfation of phenol by *p*NPS. **a)** Initial reaction rates versus phenol concentration for different *p*NPS concentrations. **b)** Double-reciprocal (Lineweaver-Burk) plot of initial reaction raters versus *p*NPS concentration at different phenol concentrations. **c)** Linear regressions of the slopes of the double-reciprocal plot as a function of phenol concentration (left) and the y-intercepts of the same plot as a function of 1/[phenol] (right). The intercept replot provides a y-intercept corresponding to 1/V_max_ and an x-intercept corresponding to −1/*K*_m,phenol_. The slope replot provides a y-intercept equal to K_m,*p*NPS_/V_max_ and an x-intercept corresponding to −*K*_i_.

### 3.3. A Two-Histidine Working Model for the Catalytic Cycle in PAPS-independent Arylsulfate sulfotransferases

The structural and kinetic observations support a catalytic cycle in which His350 links two half-reactions through a covalent intermediate (**Figure 3**). The donor first transfers its sulfo group to the enzyme and releases pNP; phenol then reacts with the sulfo-enzyme to form phenyl sulfate. This organization agrees with the established Ec_ASST_1 mechanism. His350 has a directly observed carrier function, whereas the proposed acid–base role of His283 is a mechanistic hypothesis evaluated here for the acceptor-side reaction.

**Figure 3.**
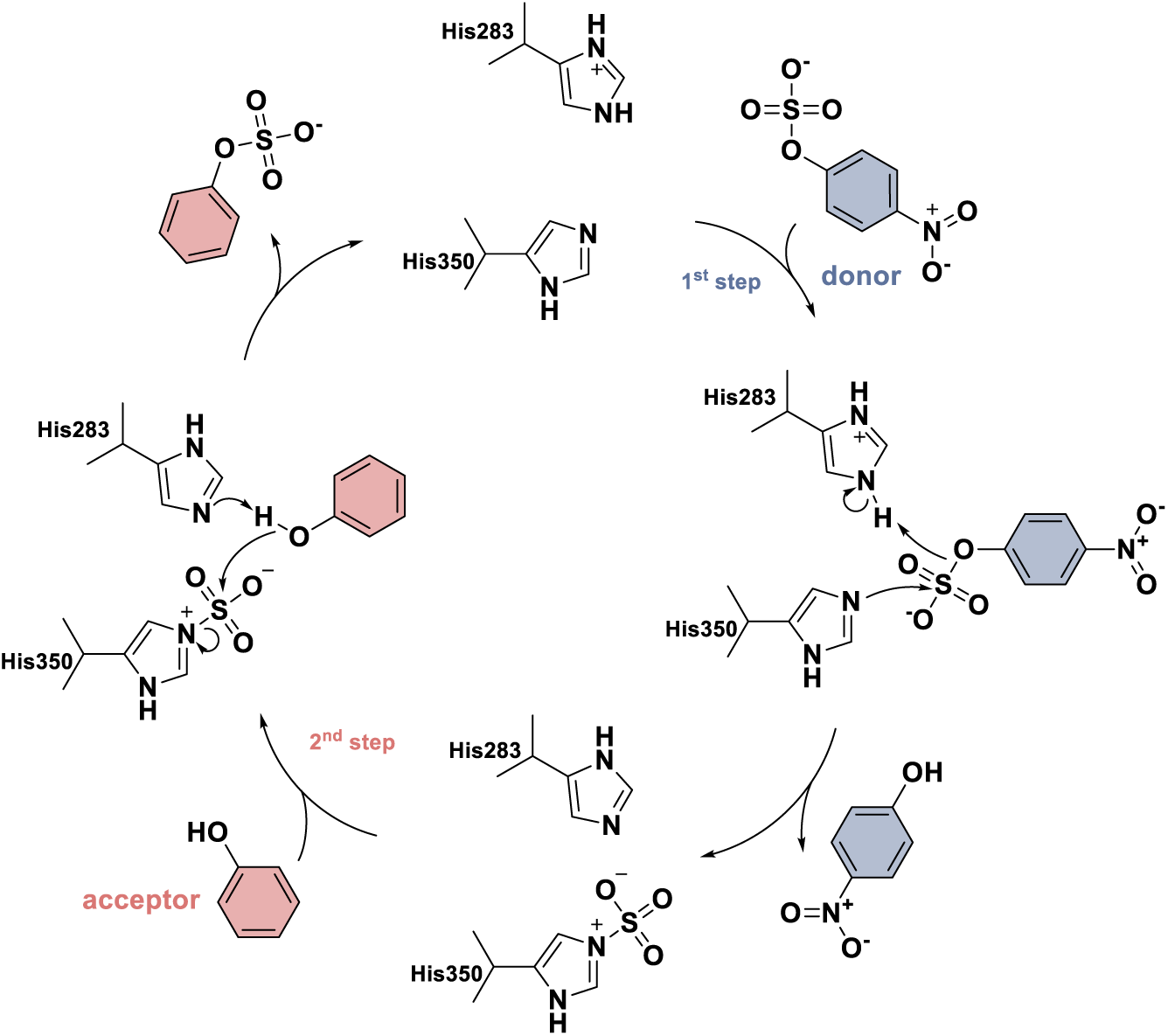
Proposed catalytic mechanism of ASST from *Desulfitobacterium hafniense* (Dh_ASST_6). The donor-side half-reaction forms covalent sulfo-His350 and releases *p*NP. In the acceptor-side half-reaction, phenol receives the sulfo group, and the unsulfated enzyme is regenerated. The scheme proposes proton donation by His283 during the first half-reaction and proton acceptance during the second. Sulfo-His350 is supported by crystallography, and the overall ping–pong organization is supported by kinetics. The first-half-reaction proton-transfer sequence was not simulated; the second-half-reaction sequence was examined using a selected neutral-His283 starting microstate.

In this cycle, His350 serves as the sulfo-carrying residue, as directly supported by the crystal structure of the sulfonated enzyme and by its correspondence to the catalytic histidine identified in Ec_ASST_1^33^. The restricted dimensions of the active-site cavity are consistent with a stepwise ping–pong mechanism, because such a compact pocket appears less compatible with simultaneous space for donor and acceptor than with alternating substrate occupancy of a substituted enzyme intermediate.

Such stepwise organization is also represented in other ping–pong bi–bi enzyme systems. Amine transaminases, for example, have been described as following a ping–pong bi–bi mechanism composed of two half-reactions, each involving several reversible chemical steps, and broader transaminase literature likewise frames catalysis as sequential donor- and acceptor-side chemistry linked by conversion of the enzyme-bound cofactor state.^93, 94^ Phosphoglycosyl transferases provide a second example, in which reaction of the first substrate generates an enzyme intermediate that is subsequently turned over in the second half-reaction.^95^

In the first half-reaction, *p*NPS transfers its sulfo group to His350, giving the sulfo-enzyme intermediate and releasing the desulfonated donor product. The active-site arrangement suggests that His283 may participate in this step. Our constant pH simulations in the absence of substrate indicate that His283 predominantly adopts a protonated state (HIP), which is consistent with a role in facilitating leaving-group departure by proton donation during sulfo-enzyme formation. In the second half-reaction, the acceptor binds to the sulfo-enzyme and receives the sulfo group from His350. His283 is proposed to assist this step by enhancing acceptor nucleophilicity and stabilizing the proton-coupled transfer geometry required for productive turnover. Our proposed mechanism is consistent with broader sulfotransferase chemistry, in which a catalytic histidine deprotonates the acceptor hydroxyl group and thereby increases its nucleophilicity for sulfo transfer; related mechanistic studies further support proton-transfer-assisted sulfuryl transfer in sulfotransferase catalysis.^96, 97^ The structural and kinetic data are complementary: the ping–pong behavior suggests formation of a substituted enzyme intermediate, whereas the crystal structure identifies that intermediate directly as sulfonated His350. Since the sulfo-enzyme state is experimentally established, but its turnover chemistry is not accessible from crystallography alone, we focused on our molecular dynamics simulations and QM/MM free-energy calculations on the second half-reaction, namely the rate limiting step, of the catalytic cycle.

### 3.4. Mechanistically Refined Simulations Capture the Reactive Sulfo-Enzyme Configuration

The model-building results reveal a distinction between structural stability and chemical readiness (**Figure 4**). Reconstruction of residues 576–590 produced 310 minimized models distributed among ten clusters (Section S3.2; **Figure S4**; **Table S4**). The largest cluster contained 72 models, or 23.2% of the ensemble, and its representative showed comparatively limited loop drift in subsequent MD. This supports its use as a plausible starting conformation. However, frequencies among generated and minimized models are not equilibrium loop populations, and the unresolved density does not establish a unique loop structure. The selected conformation is therefore a modelling choice within a heterogeneous entrance region.

**Figure 4.**
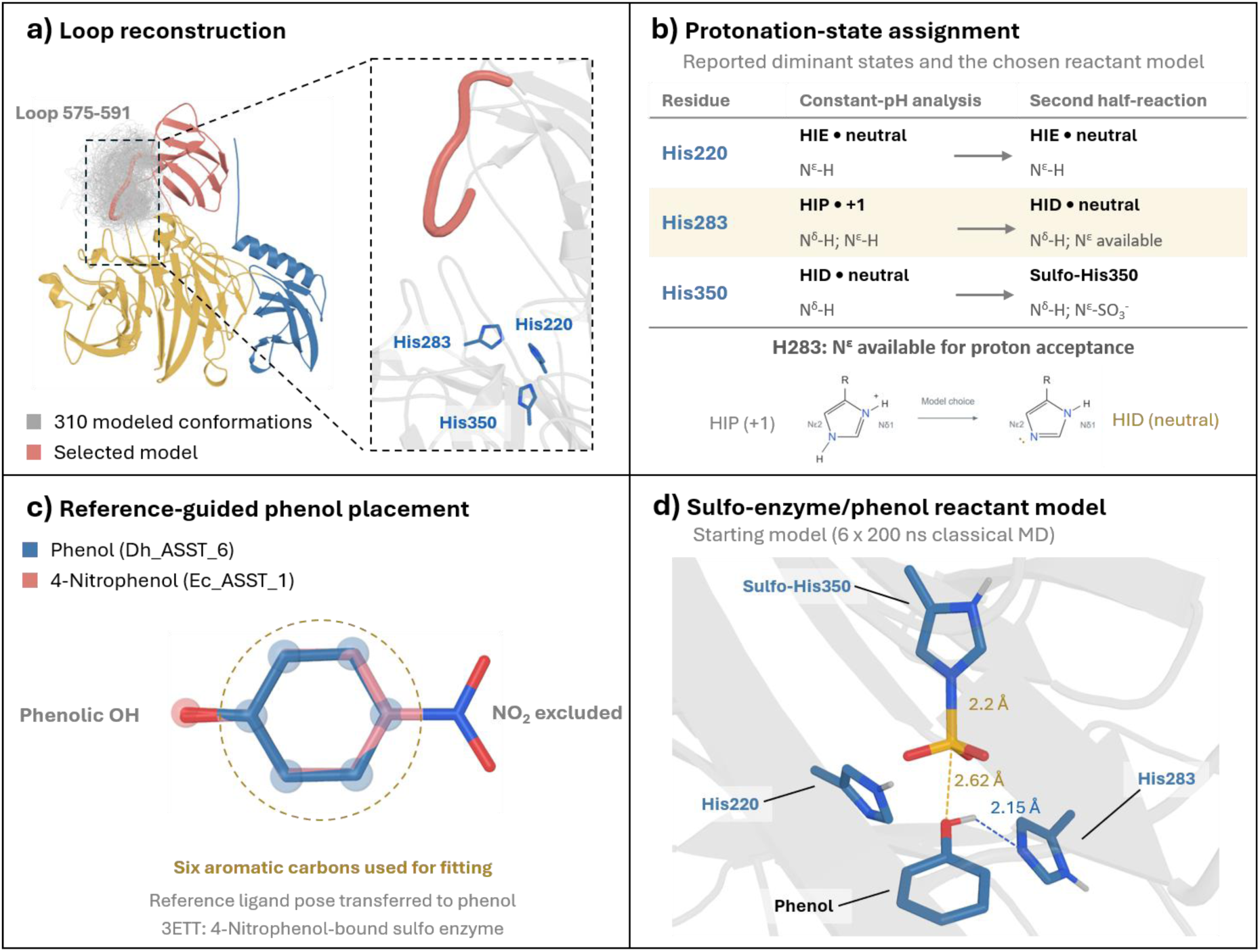
Loop reconstruction, protonation-state assignment, and preparation of the Dh_ASST_6 sulfo-enzyme/phenol complex. **a)** Ensemble of 310 sampled conformations of the unresolved loop region (residues 575–591), shown in grey, with the selected loop highlighted in coral. The close-up shows its position relative to the catalytic histidines. The N-terminal, central catalytic, and C-terminal domains are colored blue, gold, and coral, respectively. **b)** Qualitative comparison of the dominant histidine protonation states reported from constant-pH simulations and the assignments used for the second half-reaction. HIE and HID denote neutral histidine with hydrogen at Nε2 and Nδ1, respectively; HIP denotes doubly protonated histidine with a +1 charge. His220 retains the HIE state, whereas His283 is assigned HID, leaving Nε2 available for proton acceptance. His350 carries the sulfo group at Nε2 and retains hydrogen at Nδ1. The His283 schematic illustrates this model-state choice. **c)** Alignment-guided positioning of phenol (blue) using the 4-nitrophenol ligand (pink) from the Ec_ASST_1 reference structure (PDB 3ETT). Following protein alignment, the six aromatic carbon atoms were used for ligand fitting, excluding the nitro substituent. The gold outline marks the fitting region. **d)** Starting sulfo-enzyme/phenol configuration used for six independent 200 ns classical molecular dynamics simulations. The S–N bond preparation target was 2.2 Å. Dashed lines indicate the phenol O···S separation (2.62 Å) and phenol H···His283 Nε2 contact (2.15 Å) in the displayed starting model.

Protonation-state analysis introduces a more significant state choice. At pH 6.5, the reported substrate-free CpHMD trajectories predominantly sampled HIE220 (98.62%), HIP283 (100%), and HID350 (99.98%) (Section S3.4; **Table S7**). In contrast, the second-half-reaction model uses neutral HID283, with Nδ1 protonated and Nε2 available to accept the phenolic proton. Sulfo-His350 retains Nδ1–H and carries the sulfo group at Nε2. Neutral His283 is thus a mechanistically selected reactant microstate, not the dominant state predicted by the substrate-free calculation. The subsequent reaction calculation is conditional on this microstate and does not include the free-energy cost of populating it.

Phenol placement transferred the shared aromatic-ring geometry from the 4-nitrophenol-containing Ec_ASST_1 structure, PDB 3ETT, to the Dh_ASST_6 pocket (Section S3.5). This provides a homolog-informed initial pose. Across six 200 ns classical simulations, the protein backbone generally remained within approximately 1–2 Å RMSD of the starting structure, while phenol explored a broader range of active-site geometries (Section S3.6; **Figures S7– S9**). Only 3.91% of analyzed frames met both selection criteria, O···S < 5.5 Å and H···Nε2(His283) < 2.2 Å. This fraction characterizes a geometric subset of the sampled classical ensemble. It is not a measured reactive population or a kinetic commitment probability.

The limited simultaneous satisfaction of the two distance criteria suggests that acceptor positioning is an additional requirement beyond maintaining the global fold. For engineering, residues that orient the phenolic hydroxyl group may therefore matter as much as those that enlarge the pocket. This inference is consistent with the conserved catalytic center and variable surrounding architecture.

A further assumption concerns the sulfo-His350 S–N linkage, for which a 2.2 Å preparation target was used after initial tests starting at 1.6 Å did not yield the desired reaction under the steering protocol. The selected distance may influence reactant strain and the apparent barrier. Its sensitivity, together with the His283 protonation choice, must therefore be considered before assigning quantitative energetics to the experimental intermediate.

### 3.5. Steered QM/MM Trajectories Support Asynchronous Proton-Coupled Sulfo Transfer

The expanded QM region includes the reacting species and conserved neighboring side chains so that their electronic response can contribute explicitly to the modelled transformation. This is a chemical rationale for extending the minimal His283–sulfo-His350–phenol region, rather than selecting a partition because it gives a preferred barrier. Preliminary calculations show sensitivity to the QM region and steering coordinates (Section S4; **Figure S10**; **Table S8**). The production analysis comprises 314 geometrically selected starting frames, with proton-acceptor approach and O–S bond formation driven simultaneously for 2 ps. It therefore assesses a specified route through selected reactant configurations. Successful progression along that route supports mechanistic plausibility.

#### 3.5.1. Geometric Progression and the Work-Derived Profile

The Jarzynski-derived profile contains an early low-energy region followed by a larger rise as the acceptor approaches the sulfur center (**Figure 5a**). On the plotted energy scale, the six labeled configurations occur at 1.17 ps (I, −3.43 kcal mol⁻¹), 1.39 ps (II, −1.04), 1.48 ps (III, −2.13), 1.71 ps (IV, +4.52), 1.79 ps (V, +6.04), and 1.95 ps (VI, +3.63). State V lies 9.47 kcal mol⁻¹ above labeled state I, and state VI lies 7.06 kcal mol⁻¹ above I. These differences define the reference explicitly. The positive value of VI relative to the plotted zero alone is not the reaction of free-energy difference. The local extrema are features of a work-derived profile against steering time and are not independently validated transition states or metastable intermediates.

**Figure 5.**
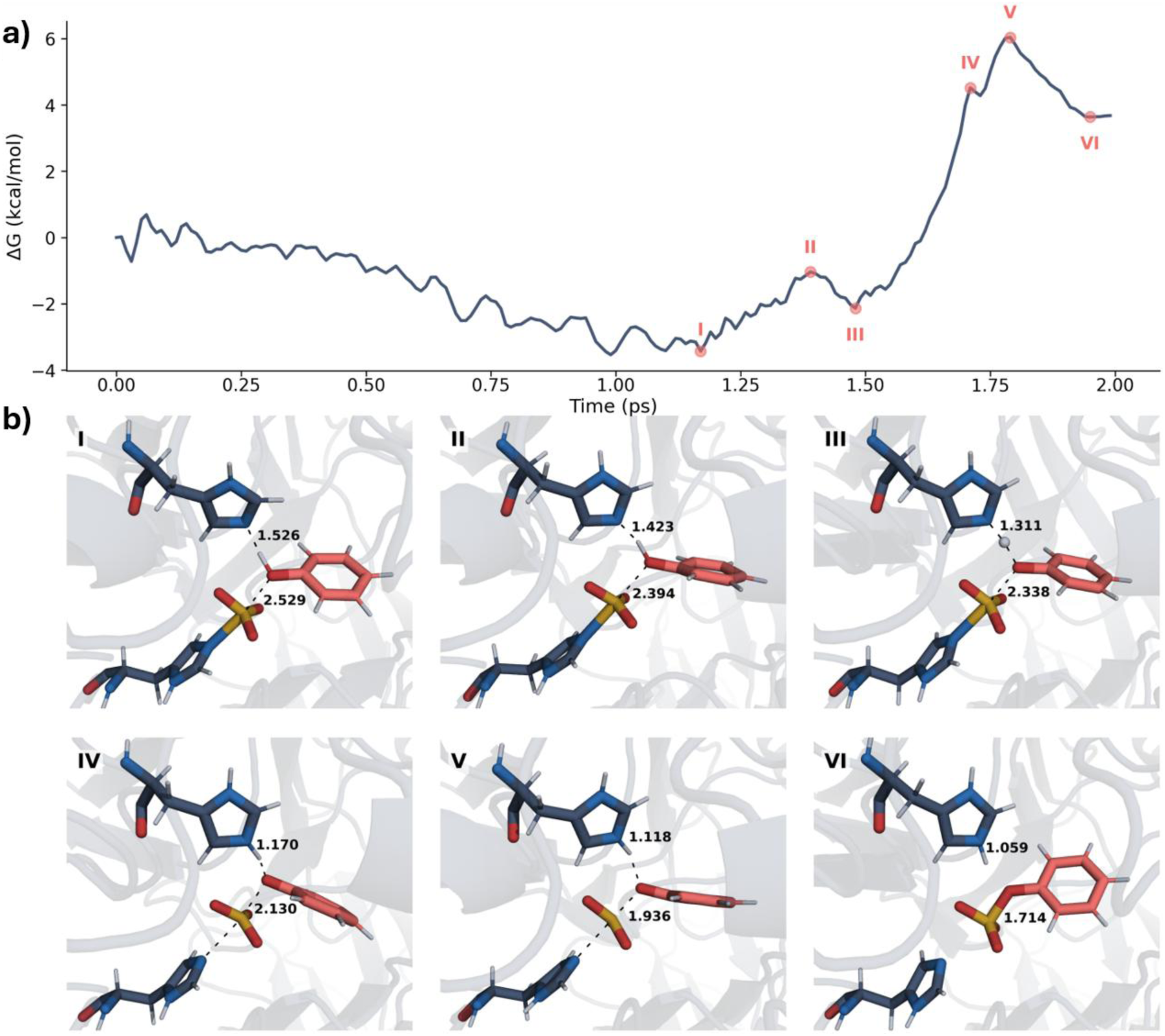
Free energy profile and representative active-site structures along the reaction pathway reconstructed from QM/MM steered molecular dynamics simulations using the Jarzynski equality. **a)** Jarzynski-derived free energy profile (ΔG, kcal mol⁻¹) as a function of simulation time. Six representative states (I–VI) are indicated, corresponding to key minima, intermediates, and high-energy regions along the reaction pathway. The system starts from a reactant-like configuration (I), passes through intermediate arrangements (II–III), and then climbs toward the main barrier region (IV–V), where the highest free energy is observed at state V, before relaxing toward the product-like state VI. b) Representative active-site snapshots corresponding to states I–VI along the Jarzynski pathway. Distances (Å) indicate the key geometric changes associated with phenol activation and sulfo-group transfer, including O_phneol_ - H_phneol_ cleavage/N_His283_ - H_His283_ formation and S_phneol_ - O_phneol_ formation/N_His350_ - S_His350_ cleavage. His350 carries the covalently bound SO_3_^-^ group, while His283 acts as the proton acceptor during the phenol activation.

Relative to the reactant minimum, the overall free-energy barrier amounts to 9.58 kcal mol⁻¹. The profile therefore indicates a two-stage barrier crossing, with a shallow intermediate separating two distinct high-energy regions. While the first barrier is comparatively small, the later part of the reaction is seen with a much steeper rise in free energy and contains the dominant energetic bottleneck. This suggests that the second catalytic step is not a single event but instead involves an initial reactive preorganization phase followed by a later bond-reorganization step. Such an observation is consistent with QM/MM analyses of transfer enzymes, which emphasize the role of active-site preorganization in shaping catalytic free-energy barriers, and with sulfotransferase-specific calculations showing that sulfuryl-transfer barriers are highly sensitive to the initial active-site conformation.^95, 98^ A related separation between early proton-transfer-related reorganization and the main sulfo-transfer barrier has been reported for human estrogen sulfotransferase, where QM/MM calculations identified proton transfer to the catalytic histidine prior to the rate-determining SO_3_-transfer event.^99^

The representative structures extracted along the profile further support this interpretation (**Figure 5b**). In the reactant region (states I–III), the system undergoes not only proton-transfer-related changes but also a clear reorientation of the acceptor within the binding pocket. In particular, the phenolic acceptor gradually rotates and shifts toward a more favorable geometry for approach to the sulfur center, while the S_His350_ - O_phneol_ distance remains too long for productive bond formation. These observations indicate that the early part of the pathway is dominated by active-site preorganization, whereas the later increase in free energy correlates with the onset of the actual bond-reorganization process. This interpretation is also consistent with QM/MM work on 3-O-sulfotransferase, in which different MD-derived starting conformations gave substantially different activation free energies, showing the importance of reactive preorganization for productive sulfuryl transfer.^98^

In the late transition region (states IV and V), proton transfer is essentially complete and the S_His350_ - O_phneol_ distance decreases further, marking progression into the sulfo-transfer event. The highest-energy configurations correspond to a highly constrained arrangement in which bond reorganization at the sulfur center takes place. Here, His283, His350, and the acceptor adopt a triangular high-energy geometry that appears to stabilize charge redistribution during bond cleavage and bond formation. In the product state (VI), the short H_phenol_ - N_His283_ distance and shortened distance confirm completion of proton transfer and formation of the new S_His350_ - O_phneol_ bond. The proposed role of His283 in this step is consistent with broader sulfotransferase chemistry, in which a catalytic histidine enhances acceptor nucleophilicity by deprotonating the acceptor hydroxyl group prior to sulfuryl transfer.^96^

This mechanistic picture is consistent with the steady-state data. The kinetic analysis identified Dh_ASST_6 as a ping–pong bi–bi enzyme, indicating formation of a substituted enzyme intermediate, whereas the crystal structure directly shows this state as sulfonated His350. The present QM/MM profile therefore provides the missing atomistic description of how this experimentally observed intermediate is converted into product. Although the calculated barrier of 9.58 kcal mol⁻¹ is not directly comparable to the apparent activation free energy derived from *k*cat, because the simulations describe only the second half-reaction from a preorganized bound reactant state, the two are qualitatively consistent in supporting a catalytically accessible turnover step. (Information about the near attack position are shown in section S5).

#### 3.5.2. CV-Space Occupancy Describes the Imposed Pathway

Projection onto CV1, the phenolic H···Nε2(His283) distance, and CV2, the phenolic O···S distance, summarizes how configurations progress under the steering protocol (**Figure 6**). The separate density maps for regions I–VI and the combined map show a narrow distribution in which both distances decrease. These maps report occupancy of the driven trajectories. Without equilibrium sampling or appropriate nonequilibrium reweighting and bias removal, high occupancy cannot be interpreted as a thermodynamically stable basin, and low occupancy cannot identify a transition state. Moreover, a narrow path is expected when two coordinates are steered together and does not exclude alternative pathways.

**Figure 6.**
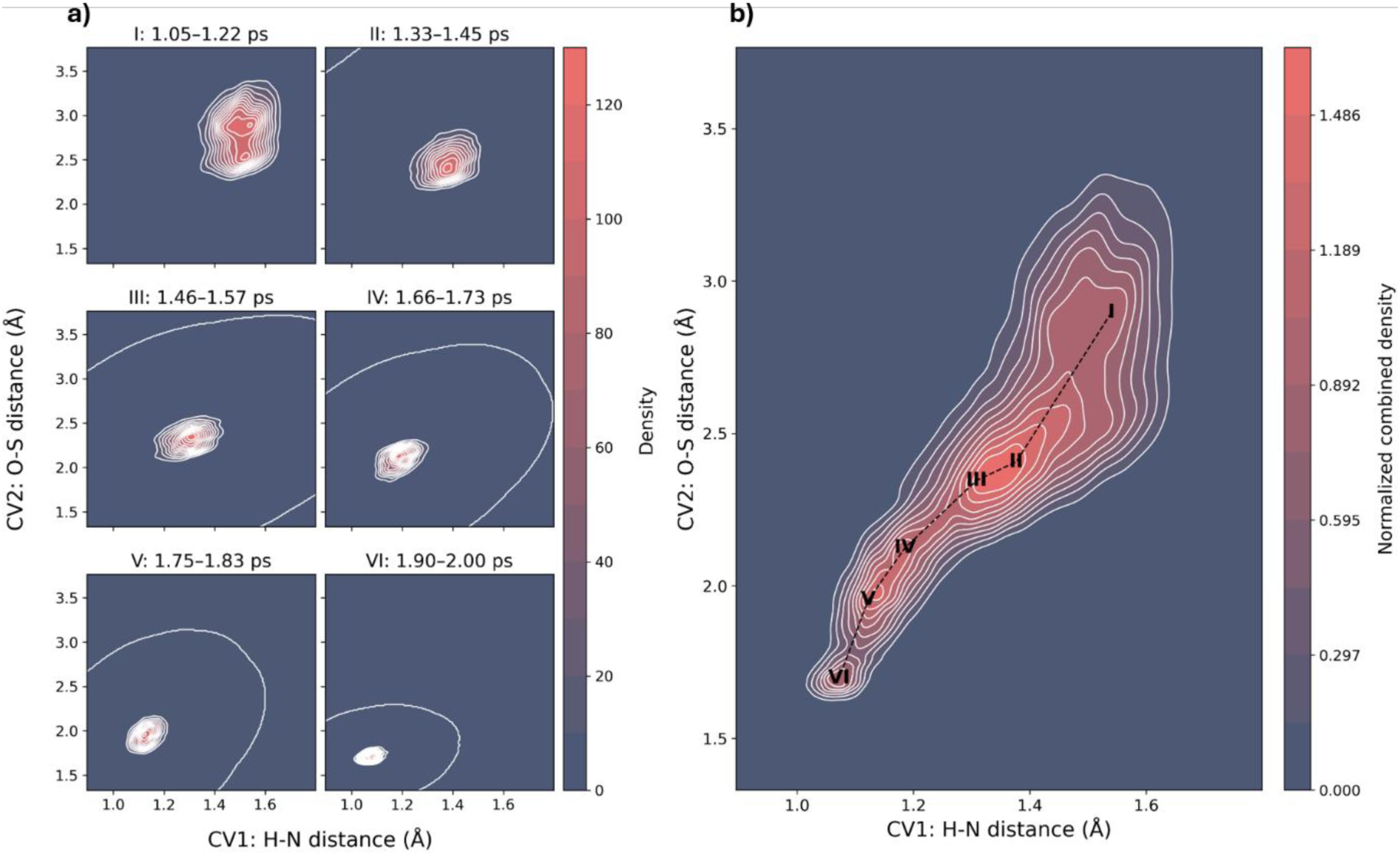
Configurational occupancy in the two steered collective variables. **a)** Kernel density estimates for configurations assigned to regions I–VI of Figure 5, projected onto CV1 [phenolic H···Nε2(His283)] and CV2 [phenolic O···S(sulfo-His350)], both in Å. **b)** Combined occupancy map with the path associated with the work-derived profile overlaid as a dashed line. Both CVs shorten during the imposed transformation. Density reflects sampling under the applied restraints and is not an equilibrium free-energy landscape. Labels I–VI denote corresponding pathway regions, not independently established stable states or transition states.

The time-colored projection confirms progression toward shorter H···N and O···S distances, and the work-colored projection associates the later part of the protocol with larger accumulated work (**Figure S13**). Together with the representative snapshots, these data support an asynchronous route in which proton transfer advances before the closest O–S approach. They do not independently demonstrate that this order is preferred under unbiased dynamics, because the pulling schedule influences both coordinates. Alternative proton-transfer partners, protonation microstates, and cleavage coordinates remain possible tests of the proposed mechanism.

#### 3.5.3. Energetic Context of the Second Half-Reaction

The calculated free-energy profile describes the acceptor-dependent half-reaction of the ASST ping–pong cycle, starting from the preformed sulfo-His350 intermediate with phenol positioned in a reactive configuration. This separation is consistent with the established catalytic mechanism of PAPS-independent ASSTs, in which sulfonation of the catalytic histidine precedes acceptor binding and subsequent sulfo transfer. Accordingly, the free-energy difference between the reactant and product basins reflects the energetics of this enzyme-bound half-reaction rather than the overall thermodynamics of sulfuryl transfer from *p*NPS to phenol. The complete catalytic cycle additionally includes sulfo-enzyme formation, donor-product release, acceptor association, and product dissociation, together with the associated conformational and protonation equilibria.

Within this mechanistic framework, the product-like state remains higher in free energy than the initial sulfo-enzyme–phenol state. Such an uphill contribution from an individual catalytic step is compatible with overall turnover because the thermodynamic driving force of the complete reaction is distributed across the two half-reactions and the associated binding and release processes. QM/MM free-energy calculations commonly focus on selected elementary steps of multistep enzymatic mechanisms, with the resulting profile describing the free-energy landscape associated with the particular reaction coordinates and states included in the simulation.^100^ The present profile is therefore most informative with respect to the activation free energy and reaction sequence, rather than as a measure of the overall equilibrium between donor and acceptor substrates.

The principal barrier of approximately 9.5 kcal mol⁻¹ is substantially lower than the apparent activation free energy derived from the experimental turnover rate. Application of transition-state theory to kcat=14.5 s⁻¹ at 39 °C gives an apparent Δ*G*^‡^ of approximately 16.6 kcal mol⁻¹. This difference is reasonable because the experimental value reflects steady-state turnover of the complete catalytic cycle, whereas the QM/MM simulations start from a preorganized, reactive sulfo-enzyme–phenol configuration and probe only the chemical acceptor half-reaction. The comparison therefore suggests that sulfo transfer itself is energetically accessible once the reactive configuration is formed and that other processes within the catalytic cycle, including substrate organization, protonation equilibria, conformational rearrangements, or product release, may contribute to the experimentally observed turnover barrier.

## 4. Conclusion

This study provided a full mechanistic picture of catalysis by the PAPS-independent sulfotransferase ASTB from *Desulfitobacterium hafniense* clade 6 (Dh_ASST_6) by integrating crystallography, steady-state kinetics, and steered QM/MM MD simulations. We determined crystal structures of Dh_ASST_6 in the native and *p*-NPS treated states at 1.9 and 2.79 Å resolution, respectively, providing direct structural insight into the conserved ASST catalytic architecture and the sulfo-enzyme state. The structural data identify His350 as the transiently sulfonated residue and define a conserved histidine-centered active site together with His283 and His220. The kinetic analysis supports a ping–pong bi–bi mechanism in which the aryl sulfate donor reacts first to generate a covalent sulfo-enzyme intermediate, followed by acceptor binding and sulfo-group transfer in the second-half reaction.

Building on this foundation, the steered QM/MM free energy simulations describe how the sulfo-enzyme intermediate is turned over at atomistic detail. The reconstructed free-energy profile shows that turnover of the sulfo-enzyme intermediate proceeds through a multistep pathway involving early active-site preorganization, proton transfer to His283, and subsequent sulfo transfer as the main bond-reorganization event. The highest-energy region is characterized by a constrained triangular arrangement of His283, His350, and the acceptor, showing how the enzyme stabilizes charge redistribution during catalysis. Structural, kinetic and QM/MM results support a mechanism in which His350 acts as the transient sulfo carrier, while His283 promotes acceptor activation and assists productive sulfo transfer.

## ASSOCIATED CONTENT

Supporting Information.

Abbreviations; supplementary crystallographic analysis and structural comparison between Dh_ASST_6 and Ec_ASST_1; detailed computational methodologies including bioinformatic analysis, loop reconstruction, tunnel analysis, structure-based docking, constant-pH simulations, and molecular dynamics simulations; evaluation of the steered QM/MM MD protocol for the second catalytic step; and characterization of near-attack conformations relevant to sulfuryl transfer.

## Author Contributions

**Conceptualization:** M.D.D., M.C., F.D.

**Experimental methodology and investigation:** N.N, H.O., D.B., L.C., M.D.D., F.D., M.C.

**Computational methodology and software:** M.D.D., D.B.

**Data curation and validation:** N.N., L.C. (Biochemistry, Enzymology, Crystallography), H.O., A.J.D. (Enzymology), D.B. (Computational Investigation)

**Formal analysis:** N.N. (Crystallography), H.O. A.J.D. (Enzymology), D.B. (Computational Investigation)

**Visualization:** D.B.

**Writing – original draft:** D.B.

**Writing – review & editing:** All authors

**Supervision:** M.C., L.C., M.D.D., F.D.

**Funding acquisition:** M.D.D., M.C., F.D.

## Notes

The authors declare that they have no conflict of interest.

## Acknowledgment

M.D.D. and D.B. thank Ahmed Ahmed Hassanin and Prof. Pedro A. Sánchez-Murcia for valuable discussions and critical scientific input.

## Declaration of Generative AI and AI-assisted technologies

During the preparation of this manuscript, the authors used ChatGPT to identify potential typographical and grammatical errors. All AI-generated suggestions and information were carefully reviewed and verified by the authors prior to incorporation into the manuscript. For language checking, prompts were limited to identifying typographical and grammatical issues without requesting modifications to the scientific content or writing style. The authors retain full responsibility for the accuracy, integrity, and final content of this publication.

## References

(1) Marques, C.; Reis, C. A.; Vives, R. R.; Magalhaes, A. Heparan Sulfate Biosynthesis and Sulfation Profiles as Modulators of Cancer Signalling and Progression. Front Oncol 2021, 11, 778752. DOI: 10.3389/fonc.2021.778752.

(2) Kurogi, K.; Rasool, M. I.; Alherz, F. A.; El Daibani, A. A.; Bairam, A. F.; Abunnaja, M. S.; Yasuda, S.; Wilson, L. J.; Hui, Y.; Liu, M. C. SULT genetic polymorphisms: physiological, pharmacological and clinical implications. Expert Opin Drug Metab Toxicol 2021, 17 (7), 767–784. DOI: 10.1080/17425255.2021.1940952.

(3) Kauffman, F. C. Sulfonation in pharmacology and toxicology. Drug Metab Rev 2004, 36 (3-4), 823–843. DOI: 10.1081/dmr-200033496.

(4) Hempel, N.; Gamage, N.; Martin, J. L.; McManus, M. E. Human cytosolic sulfotransferase SULT1A1. Int J Biochem Cell Biol 2007, 39 (4), 685–689. DOI: 10.1016/j.biocel.2006.10.002.

(5) Suiko, M.; Kurogi, K.; Hashiguchi, T.; Sakakibara, Y.; Liu, M. C. Updated perspectives on the cytosolic sulfotransferases (SULTs) and SULT-mediated sulfation. Biosci Biotechnol Biochem 2017, 81 (1), 63–72. DOI: 10.1080/09168451.2016.1222266.

(6) Pedersen, L. C.; Yi, M.; Pedersen, L. G.; Kaminski, A. M. From Steroid and Drug Metabolism to Glycobiology, Using Sulfotransferase Structures to Understand and Tailor Function. Drug Metab Dispos 2022, 50 (7), 1027–1041. DOI: 10.1124/dmd.121.000478.

(7) Monterrey, D. T.; Benito-Arenas, R.; Revuelta, J.; Garcia-Junceda, E. Design of a biocatalytic cascade for the enzymatic sulfation of unsulfated chondroitin with in situ generation of PAPS. Front Bioeng Biotechnol 2023, 11, 1099924. DOI: 10.3389/fbioe.2023.1099924.

(8) Gamage, N.; Barnett, A.; Hempel, N.; Duggleby, R. G.; Windmill, K. F.; Martin, J. L.; McManus, M. E. Human sulfotransferases and their role in chemical metabolism. Toxicol Sci 2006, 90 (1), 5–22. DOI: 10.1093/toxsci/kfj061.

(9) Mueller, J. W.; Gilligan, L. C.; Idkowiak, J.; Arlt, W.; Foster, P. A. The Regulation of Steroid Action by Sulfation and Desulfation. Endocr Rev 2015, 36 (5), 526–563. DOI: 10.1210/er.2015-1036.

(10) Langford, L.; Shah, D. D. Bioinformatic Analysis of Sulfotransferases from an Unexplored Gut Microbe, Sutterella wadsworthensis 3_1_45B: Possible Roles towards Detoxification via Sulfonation by Members of the Human Gut Microbiome. Int J Mol Sci 2024, 25 (5). DOI: 10.3390/ijms25052983.

(11) Coughtrie, M. W.; Sharp, S.; Maxwell, K.; Innes, N. P. Biology and function of the reversible sulfation pathway catalysed by human sulfotransferases and sulfatases. Chem Biol Interact 1998, 109 (1-3), 3–27. DOI: 10.1016/s0009-2797(97)00117-8.

(12) Runge-Morris, M.; Kocarek, T. A. Regulation of sulfotransferase and UDP-glucuronosyltransferase gene expression by the PPARs. PPAR Res 2009, 2009, 728941. DOI: 10.1155/2009/728941.

(13) Paritala, H.; Carroll, K. S. New targets and inhibitors of mycobacterial sulfur metabolism. Infect Disord Drug Targets 2013, 13 (2), 85–115. DOI: 10.2174/18715265113139990022.

(14) van der Horst, M. A.; Hartog, A. F.; El Morabet, R.; Marais, A.; Kircz, M.; Wever, R. Enzymatic Sulfation of Phenolic Hydroxy Groups of Various Plant Metabolites by an Arylsulfotransferase. European Journal of Organic Chemistry 2014, 2015 (3), 534–541. DOI: 10.1002/ejoc.201402875.

(15) Marhol, P.; Hartog, A. F.; van der Horst, M. A.; Wever, R.; Purchartová, K.; Fuksová, K.; Kuzma, M.; Cvačka, J.; Křen, V. Preparation of silybin and isosilybin sulfates by sulfotransferase from Desulfitobacterium hafniense. Journal of Molecular Catalysis B: Enzymatic 2013, 89, 24–27. DOI: 10.1016/j.molcatb.2012.12.005.

(16) Roubalova, L.; Purchartova, K.; Papouskova, B.; Vacek, J.; Kren, V.; Ulrichova, J.; Vrba, J. Sulfation modulates the cell uptake, antiradical activity and biological effects of flavonoids in vitro: An examination of quercetin, isoquercitrin and taxifolin. Bioorg Med Chem 2015, 23 (17), 5402–5409. DOI: 10.1016/j.bmc.2015.07.055.

(17) Kim, D. H.; Kim, H. S.; Kobashi, K. Purification and characterization of novel sulfotransferase obtained from Klebsiella K-36, an intestinal bacterium of rat. J Biochem 1992, 112 (4), 456–460. DOI: 10.1093/oxfordjournals.jbchem.a123921.

(18) van der Horst, M. A.; van Lieshout, J. F. T.; Bury, A.; Hartog, A. F.; Wever, R. Sulfation of Various Alcoholic Groups by an Arylsulfate Sulfotransferase from Desulfitobacterium hafniense and Synthesis of Estradiol Sulfate. Advanced Synthesis & Catalysis 2012, 354 (18), 3501–3508. DOI: 10.1002/adsc.201200564.

(19) Stone, M. J.; Chuang, S.; Hou, X.; Shoham, M.; Zhu, J. Z. Tyrosine sulfation: an increasingly recognised post-translational modification of secreted proteins. N Biotechnol 2009, 25 (5), 299–317. DOI: 10.1016/j.nbt.2009.03.011.

(20) Muthana, S. M.; Campbell, C. T.; Gildersleeve, J. C. Modifications of glycans: biological significance and therapeutic opportunities. ACS Chem Biol 2012, 7 (1), 31–43. DOI: 10.1021/cb2004466.

(21) Stewart, V.; Ronald, P. C. Sulfotyrosine residues: Interaction specificity determinants for extracellular protein-protein interactions. J Biol Chem 2022, 298 (8), 102232. DOI: 10.1016/j.jbc.2022.102232.

(22) Al-Horani, R. A.; Desai, U. R. Chemical Sulfation of Small Molecules - Advances and Challenges. Tetrahedron 2010, 66 (16), 2907–2918. DOI: 10.1016/j.tet.2010.02.015.

(23) Sheldon, R. A.; Brady, D.; Bode, M. L. The Hitchhiker’s guide to biocatalysis: recent advances in the use of enzymes in organic synthesis. Chem Sci 2020, 11 (10), 2587–2605. DOI: 10.1039/c9sc05746c.

(24) Esposito, F.; Laezza, A.; Gargiulo, V.; Traboni, S.; Iadonisi, A.; La Gatta, A.; Schiraldi, C.; Bedini, E. Multi-step Strategies Toward Regioselectively Sulfated M-Rich Alginates. Biomacromolecules 2023, 24 (6), 2522–2531. DOI: 10.1021/acs.biomac.3c00045.

(25) Negishi, M.; Pedersen, L. G.; Petrotchenko, E.; Shevtsov, S.; Gorokhov, A.; Kakuta, Y.; Pedersen, L. C. Structure and function of sulfotransferases. Arch Biochem Biophys 2001, 390 (2), 149–157. DOI: 10.1006/abbi.2001.2368.

(26) Badri, A.; Williams, A.; Awofiranye, A.; Datta, P.; Xia, K.; He, W.; Fraser, K.; Dordick, J. S.; Linhardt, R. J.; Koffas, M. A. G. Complete biosynthesis of a sulfated chondroitin in Escherichia coli. Nat Commun 2021, 12 (1), 1389. DOI: 10.1038/s41467-021-21692-5.

(27) Hall, M. Enzymatic strategies for asymmetric synthesis. RSC Chem Biol 2021, 2 (4), 958–989. DOI: 10.1039/d1cb00080b.

(28) Kissman, E. N.; Sosa, M. B.; Millar, D. C.; Koleski, E. J.; Thevasundaram, K.; Chang, M. C. Y. Expanding chemistry through in vitro and in vivo biocatalysis. Nature 2024, 631 (8019), 37–48. DOI: 10.1038/s41586-024-07506-w.

(29) Winkler, C. K.; Schrittwieser, J. H.; Kroutil, W. Power of Biocatalysis for Organic Synthesis. ACS Cent Sci 2021, 7 (1), 55–71. DOI: 10.1021/acscentsci.0c01496.

(30) Ji, Y.; Islam, S.; Cui, H.; Dhoke, G. V.; Davari, M. D.; Mertens, A. M.; Schwaneberg, U. Loop engineering of aryl sulfotransferase B for improving catalytic performance in regioselective sulfation. Catalysis Science & Technology 2020, 10 (8), 2369–2377. DOI: 10.1039/d0cy00063a.

(31) Chapman, E.; Best, M. D.; Hanson, S. R.; Wong, C. H. Sulfotransferases: structure, mechanism, biological activity, inhibition, and synthetic utility. Angew Chem Int Ed Engl 2004, 43 (27), 3526–3548. DOI: 10.1002/anie.200300631.

(32) Malojcic, G.; Glockshuber, R. The PAPS-independent aryl sulfotransferase and the alternative disulfide bond formation system in pathogenic bacteria. Antioxid Redox Signal 2010, 13 (8), 1247– 1259. DOI: 10.1089/ars.2010.3119.

(33) Malojcic, G.; Owen, R. L.; Grimshaw, J. P.; Brozzo, M. S.; Dreher-Teo, H.; Glockshuber, R. A structural and biochemical basis for PAPS-independent sulfuryl transfer by aryl sulfotransferase from uropathogenic Escherichia coli. Proc Natl Acad Sci U S A 2008, 105 (49), 19217–19222. DOI: 10.1073/pnas.0806997105.

(34) Bojarova, P.; Williams, S. J. Sulfotransferases, sulfatases and formylglycine-generating enzymes: a sulfation fascination. Curr Opin Chem Biol 2008, 12 (5), 573–581. DOI: 10.1016/j.cbpa.2008.06.018.

(35) Zhao, H.; van der Donk, W. A. Regeneration of cofactors for use in biocatalysis. Curr Opin Biotechnol 2003, 14 (6), 583–589. DOI: 10.1016/j.copbio.2003.09.007.

(36) Islam, S.; Mate, D. M.; Martinez, R.; Jakob, F.; Schwaneberg, U. A robust protocol for directed aryl sulfotransferase evolution toward the carbohydrate building block GlcNAc. Biotechnol Bioeng 2018, 115 (5), 1106–1115. DOI: 10.1002/bit.26535.

(37) Kobashi, K.; Kim, D.-H.; Morikawa, T. A novel type of arylsulfotransferase. Journal of Protein Chemistry 1987, 6 (3), 237–244. DOI: 10.1007/bf00250287.

(38) Funabashi, M.; Baba, S.; Nonaka, K.; Hosobuchi, M.; Fujita, Y.; Shibata, T.; Van Lanen, S. G. The biosynthesis of liposidomycin-like A-90289 antibiotics featuring a new type of sulfotransferase. Chembiochem 2010, 11 (2), 184–190. DOI: 10.1002/cbic.200900665.

(39) Hartog, A. F.; Wever, R. Sulfation made easy: A new versatile donor for enzymatic sulfation by a bacterial arylsulfotransferase. Journal of Molecular Catalysis B: Enzymatic 2016, 129, 43–46. DOI: 10.1016/j.molcatb.2016.04.007.

(40) Kaysser, L.; Eitel, K.; Tanino, T.; Siebenberg, S.; Matsuda, A.; Ichikawa, S.; Gust, B. A new arylsulfate sulfotransferase involved in liponucleoside antibiotic biosynthesis in streptomycetes. J Biol Chem 2010, 285 (17), 12684–12694. DOI: 10.1074/jbc.M109.094490.

(41) Islam, S.; Laaf, D.; Infanzon, B.; Pelantova, H.; Davari, M. D.; Jakob, F.; Kren, V.; Elling, L.; Schwaneberg, U. KnowVolution Campaign of an Aryl Sulfotransferase Increases Activity toward Cellobiose. Chemistry 2018, 24 (64), 17117–17124. DOI: 10.1002/chem.201803729.

(42) Brodsky, K.; Petrankova, B.; Petraskova, L.; Pelantova, H.; Kren, V.; Valentova, K.; Bojarova, P. New Bacterial Aryl Sulfotransferases: Effective Tools for Sulfation of Polyphenols. J Agric Food Chem 2024, 72 (40), 22208–22216. DOI: 10.1021/acs.jafc.4c06771.

(43) Baek, M. C.; Kwon, A. R.; Chung, Y. J.; Kim, B. K.; Choi, E. C. Distribution of bacteria with the arylsulfate sulfotransferase activity. Arch Pharm Res 1998, 21 (4), 475–477. DOI: 10.1007/BF02974646.

(44) Malojcic, G.; Owen, R. L.; Glockshuber, R. Structural and mechanistic insights into the PAPS-independent sulfotransfer catalyzed by bacterial aryl sulfotransferase and the role of the DsbL/Dsbl system in its folding. Biochemistry 2014, 53 (11), 1870–1877. DOI: 10.1021/bi401725j.

(45) Kwon, A. R.; Yun, H. J.; Choi, E. C. Kinetic mechanism and identification of the active site tyrosine residue in Enterobacter amnigenus arylsulfate sulfotransferase. Biochem Biophys Res Commun 2001, 285 (2), 526–529. DOI: 10.1006/bbrc.2001.5185.

(46) Lee, N. S.; Kim, B. T.; Kim, D. H.; Kobashi, K. Purification and reaction mechanism of arylsulfate sulfotransferase from Haemophilus K-12, a mouse intestinal bacterium. J Biochem 1995, 118 (4), 796– 801. DOI: 10.1093/oxfordjournals.jbchem.a124982.

(47) Brodsky, K.; Kanova, K.; Konvalinkova, D.; Slamova, K.; Pelantova, H.; Valentova, K.; Bojarova, P.; Kren, V.; Petraskova, L. Bacterial Aryl Sulfotransferases in Selective and Sustainable Sulfation of Biologically Active Compounds using Novel Sulfate Donors. ChemSusChem 2022, 15 (18), e202201253. DOI: 10.1002/cssc.202201253.

(48) Oroz-Guinea, I.; Rath, M.; Tischler, I.; Ditrich, K.; Schachtschabel, D.; Breuer, M.; Kroutil, W. Biocatalytic sulfation of aromatic and aliphatic alcohols catalyzed by arylsulfate sulfotransferases. Appl Microbiol Biotechnol 2024, 108 (1), 520. DOI: 10.1007/s00253-024-13354-5.

(49) Kanova, K.; Petraskova, L.; Pelantova, H.; Rybkova, Z.; Malachova, K.; Cvacka, J.; Kren, V.; Valentova, K. Sulfated Metabolites of Luteolin, Myricetin, and Ampelopsin: Chemoenzymatic Preparation and Biophysical Properties. J Agric Food Chem 2020, 68 (40), 11197–11206. DOI: 10.1021/acs.jafc.0c03997.

(50) Valentova, K.; Purchartova, K.; Rydlova, L.; Roubalova, L.; Biedermann, D.; Petraskova, L.; Krenkova, A.; Pelantova, H.; Holeckova-Moravcova, V.; Tesarova, E.;, et al. Sulfated Metabolites of Flavonolignans and 2,3-Dehydroflavonolignans: Preparation and Properties. Int J Mol Sci 2018, 19 (8). DOI: 10.3390/ijms19082349.

(51) Purchartová, K.; Valentová, K.; Pelantová, H.; Marhol, P.; Cvačka, J.; Havlíček, L.; Křenková, A.; Vavříková, E.; Biedermann, D.; Chambers, C. S.;, et al. Prokaryotic and Eukaryotic Aryl Sulfotransferases: Sulfation of Quercetin and Its Derivatives. ChemCatChem 2015, 7 (19), 3152–3162. DOI: 10.1002/cctc.201500298.

(52) Begines, P.; Biedermann, D.; Valentova, K.; Petraskova, L.; Pelantova, H.; Maya, I.; Fernandez-Bolanos, J. G.; Kren, V. Chemoenzymatic Synthesis and Radical Scavenging of Sulfated Hydroxytyrosol, Tyrosol, and Acetylated Derivatives. J Agric Food Chem 2019, 67 (26), 7281–7288. DOI: 10.1021/acs.jafc.9b01065.

(53) Kabsch, W. Xds. Acta Crystallogr D Biol Crystallogr 2010, 66 (Pt 2), 125–132. DOI: 10.1107/S0907444909047337.

(54) Winn, M. D.; Ballard, C. C.; Cowtan, K. D.; Dodson, E. J.; Emsley, P.; Evans, P. R.; Keegan, R. M.; Krissinel, E. B.; Leslie, A. G.; McCoy, A.;, et al. Overview of the CCP4 suite and current developments. Acta Crystallogr D Biol Crystallogr 2011, 67 (Pt 4), 235–242. DOI: 10.1107/S0907444910045749.

(55) McCoy, A. J.; Grosse-Kunstleve, R. W.; Adams, P. D.; Winn, M. D.; Storoni, L. C.; Read, R. J. Phaser crystallographic software. J Appl Crystallogr 2007, 40 (Pt 4), 658–674. DOI: 10.1107/S0021889807021206.

(56) Mirdita, M.; Schutze, K.; Moriwaki, Y.; Heo, L.; Ovchinnikov, S.; Steinegger, M. ColabFold: making protein folding accessible to all. Nat Methods 2022, 19 (6), 679–682. DOI: 10.1038/s41592-022-01488-1.

(57) Murshudov, G. N.; Skubak, P.; Lebedev, A. A.; Pannu, N. S.; Steiner, R. A.; Nicholls, R. A.; Winn, M. D.; Long, F.; Vagin, A. A. REFMAC5 for the refinement of macromolecular crystal structures. Acta Crystallogr D Biol Crystallogr 2011, 67 (Pt 4), 355–367. DOI: 10.1107/S0907444911001314.

(58) Emsley, P.; Lohkamp, B.; Scott, W. G.; Cowtan, K. Features and development of Coot. Acta Crystallogr D Biol Crystallogr 2010, 66 (Pt 4), 486–501. DOI: 10.1107/S0907444910007493.

(59) Chen, V. B.; Arendall, W. B., 3rd; Headd, J. J.; Keedy, D. A.; Immormino, R. M.; Kapral, G. J.; Murray, L. W.; Richardson, J. S.; Richardson, D. C. MolProbity: all-atom structure validation for macromolecular crystallography. Acta Crystallogr D Biol Crystallogr 2010, 66 (Pt 1), 12–21. DOI: 10.1107/S0907444909042073.

(60) Krieger, E.; Vriend, G. YASARA View - molecular graphics for all devices - from smartphones to workstations. Bioinformatics 2014, 30 (20), 2981–2982. DOI: 10.1093/bioinformatics/btu426.

(61) Schrodinger, LLC. The PyMOL Molecular Graphics System, Version 1.8. 2015.

(62) Lua, R. C.; Wilson, S. J.; Konecki, D. M.; Wilkins, A. D.; Venner, E.; Morgan, D. H.; Lichtarge, O. UET: a database of evolutionarily-predicted functional determinants of protein sequences that cluster as functional sites in protein structures. Nucleic Acids Res 2016, 44 (D1), D308–312. DOI: 10.1093/nar/gkv1279.

(63) Camacho, C.; Coulouris, G.; Avagyan, V.; Ma, N.; Papadopoulos, J.; Bealer, K.; Madden, T. L. BLAST+: architecture and applications. BMC Bioinformatics 2009, 10, 421. DOI: 10.1186/1471-2105-10-421.

(64) Information, N. C. f. B. NCBI Multiple Sequence Alignment Viewer. U.S. National Library of Medicine, 2026. https://www.ncbi.nlm.nih.gov/projects/msaviewer/ (accessed April 30, 2026).

(65) Chovancova, E.; Pavelka, A.; Benes, P.; Strnad, O.; Brezovsky, J.; Kozlikova, B.; Gora, A.; Sustr, V.; Klvana, M.; Medek, P.;, et al. CAVER 3.0: a tool for the analysis of transport pathways in dynamic protein structures. PLoS Comput Biol 2012, 8 (10), e1002708. DOI: 10.1371/journal.pcbi.1002708.

(66) Guerra, J. V. S.; Ribeiro-Filho, H. V.; Pereira, J. G. C.; Lopes-de-Oliveira, P. S. KVFinder-web: a web-based application for detecting and characterizing biomolecular cavities. Nucleic Acids Res 2023, 51 (W1), W289–W297. DOI: 10.1093/nar/gkad324.

(67) Mongan, J.; Case, D. A.; McCammon, J. A. Constant pH molecular dynamics in generalized Born implicit solvent. J Comput Chem 2004, 25 (16), 2038–2048. DOI: 10.1002/jcc.20139.

(68) Case, D. A.; Aktulga, H. M.; Belfon, K.; Cerutti, D. S.; Cisneros, G. A.; Cruzeiro, V. W. D.; Forouzesh, N.; Giese, T. J.; Gotz, A. W.; Gohlke, H.;, et al. AmberTools. J Chem Inf Model 2023, 63 (20), 6183–6191. DOI: 10.1021/acs.jcim.3c01153.

(69) Ryckaert, J.-P.; Ciccotti, G.; Berendsen, H. J. C. Numerical integration of the cartesian equations of motion of a system with constraints: molecular dynamics of n-alkanes. Journal of Computational Physics 1977, 23 (3), 327–341. DOI: 10.1016/0021-9991(77)90098-5.

(70) Anandakrishnan, R.; Aguilar, B.; Onufriev, A. V. H++ 3.0: automating pK prediction and the preparation of biomolecular structures for atomistic molecular modeling and simulations. Nucleic Acids Res 2012, 40 (Web Server issue), W537–541. DOI: 10.1093/nar/gks375.

(71) Tian, C.; Kasavajhala, K.; Belfon, K. A. A.; Raguette, L.; Huang, H.; Migues, A. N.; Bickel, J.; Wang, Y.; Pincay, J.; Wu, Q.;, et al. ff19SB: Amino-Acid-Specific Protein Backbone Parameters Trained against Quantum Mechanics Energy Surfaces in Solution. J Chem Theory Comput 2020, 16 (1), 528–552. DOI: 10.1021/acs.jctc.9b00591.

(72) Wang, J.; Wang, W.; Kollman, P. A.; Case, D. A. Automatic atom type and bond type perception in molecular mechanical calculations. J Mol Graph Model 2006, 25 (2), 247–260. DOI: 10.1016/j.jmgm.2005.12.005.

(73) Becke, A. D. Density-functional thermochemistry. III. The role of exact exchange. The Journal of Chemical Physics 1993, 98 (7), 5648–5652. DOI: 10.1063/1.464913.

(74) Lee, C.; Yang, W.; Parr, R. G. Development of the Colle-Salvetti correlation-energy formula into a functional of the electron density. Phys Rev B Condens Matter 1988, 37 (2), 785–789. DOI: 10.1103/physrevb.37.785.

(75) Hehre, W. J.; Ditchfield, R.; Pople, J. A. Self—Consistent Molecular Orbital Methods. XII. Further Extensions of Gaussian—Type Basis Sets for Use in Molecular Orbital Studies of Organic Molecules. The Journal of Chemical Physics 1972, 56 (5), 2257–2261. DOI: 10.1063/1.1677527.

(76) Hariharan, P. C.; Pople, J. A. The influence of polarization functions on molecular orbital hydrogenation energies. Theoretica Chimica Acta 1973, 28 (3), 213–222. DOI: 10.1007/bf00533485.

(77) Clark, T.; Chandrasekhar, J.; Spitznagel, G. W.; Schleyer, P. V. R. Efficient diffuse function- augmented basis sets for anion calculations. III. The 3-21+G basis set for first-row elements, Li–F. Journal of Computational Chemistry 2004, 4 (3), 294–301. DOI: 10.1002/jcc.540040303.

(78) Gaussian 16 Rev. C.01; Wallingford, CT, 2016.

(79) Bayly, C. I.; Cieplak, P.; Cornell, W.; Kollman, P. A. A well-behaved electrostatic potential based method using charge restraints for deriving atomic charges: the RESP model. The Journal of Physical Chemistry 2002, 97 (40), 10269–10280. DOI: 10.1021/j100142a004.

(80) Roothaan, C. C. J. New Developments in Molecular Orbital Theory. Reviews of Modern Physics 1951, 23 (2), 69–89. DOI: 10.1103/RevModPhys.23.69.

(81) Izadi, S.; Anandakrishnan, R.; Onufriev, A. V. Building Water Models: A Different Approach. J Phys Chem Lett 2014, 5 (21), 3863–3871. DOI: 10.1021/jz501780a.

(82) Roe, D. R.; Cheatham, T. E., 3rd. PTRAJ and CPPTRAJ: Software for Processing and Analysis of Molecular Dynamics Trajectory Data. J Chem Theory Comput 2013, 9 (7), 3084–3095. DOI: 10.1021/ct400341p.

(83) Michaud-Agrawal, N.; Denning, E. J.; Woolf, T. B.; Beckstein, O. MDAnalysis: a toolkit for the analysis of molecular dynamics simulations. J Comput Chem 2011, 32 (10), 2319–2327. DOI: 10.1002/jcc.21787.

(84) Field, M. J.; Bash, P. A.; Karplus, M. A combined quantum mechanical and molecular mechanical potential for molecular dynamics simulations. Journal of Computational Chemistry 2004, 11 (6), 700–733. DOI: 10.1002/jcc.540110605.

(85) Gaus, M.; Cui, Q.; Elstner, M. DFTB3: Extension of the self-consistent-charge density-functional tight-binding method (SCC-DFTB). J Chem Theory Comput 2012, 7 (4), 931–948. DOI: 10.1021/ct100684s.

(86) Gaus, M.; Goez, A.; Elstner, M. Parametrization and Benchmark of DFTB3 for Organic Molecules. J Chem Theory Comput 2013, 9 (1), 338–354. DOI: 10.1021/ct300849w.

(87) Gaus, M.; Lu, X.; Elstner, M.; Cui, Q. Parameterization of DFTB3/3OB for Sulfur and Phosphorus for Chemical and Biological Applications. J Chem Theory Comput 2014, 10 (4), 1518–1537. DOI: 10.1021/ct401002w.

(88) Roston, D.; Lu, X.; Fang, D.; Demapan, D.; Cui, Q. Analysis of Phosphoryl-Transfer Enzymes with QM/MM Free Energy Simulations. Methods Enzymol 2018, 607, 53–90. DOI: 10.1016/bs.mie.2018.05.005.

(89) Wang, X.; Li, R.; Cui, W.; Li, Q.; Yao, J. QM/MM free energy Simulations of an efficient Gluten Hydrolase (Kuma030) Implicate for a Reactant-State Based Protein-Design Strategy for General Acid/Base Catalysis. Sci Rep 2018, 8 (1), 7042. DOI: 10.1038/s41598-018-25471-z.

(90) Jarzynski, C. Nonequilibrium Equality for Free Energy Differences. Physical Review Letters 1997, 78 (14), 2690–2693. DOI: 10.1103/PhysRevLett.78.2690.

(91) Segel, I. H. Enzyme Kinetics: Behavior and Analysis of Rapid Equilibrium and Steady-State Enzyme Systems; John Wiley & Sons, 1993.

(92) Copeland, R. A. Enzymes: A Practical Introduction to Structure, Mechanism, and Data Analysis; John Wiley & Sons, 2023. DOI: 10.1002/9781119793304.

(93) Koper, K.; Han, S. W.; Pastor, D. C.; Yoshikuni, Y.; Maeda, H. A. Evolutionary origin and functional diversification of aminotransferases. J Biol Chem 2022, 298 (8), 102122. DOI: 10.1016/j.jbc.2022.102122.

(94) Meng, Q.; Ramírez-Palacios, C.; Wijma, H. J.; Janssen, D. B. Protein engineering of amine transaminases. Frontiers in Catalysis 2022, 2. DOI: 10.3389/fctls.2022.1049179.

(95) Das, D.; Kuzmic, P.; Imperiali, B. Analysis of a dual domain phosphoglycosyl transferase reveals a ping-pong mechanism with a covalent enzyme intermediate. Proc Natl Acad Sci U S A 2017, 114 (27), 7019–7024. DOI: 10.1073/pnas.1703397114.

(96) Pedersen, L. C.; Petrotchenko, E.; Shevtsov, S.; Negishi, M. Crystal structure of the human estrogen sulfotransferase-PAPS complex: evidence for catalytic role of Ser137 in the sulfuryl transfer reaction. J Biol Chem 2002, 277 (20), 17928–17932. DOI: 10.1074/jbc.M111651200.

(97) Zhang, H.; Varlamova, O.; Vargas, F. M.; Falany, C. N.; Leyh, T. S. Sulfuryl transfer: the catalytic mechanism of human estrogen sulfotransferase. J Biol Chem 1998, 273 (18), 10888–10892. DOI: 10.1074/jbc.273.18.10888.

(98) Sousa, R. P.; Fernandes, P. A.; Ramos, M. J.; Bras, N. F. Insights into the reaction mechanism of 3-O-sulfotransferase through QM/MM calculations. Phys Chem Chem Phys 2016, 18 (16), 11488–11496. DOI: 10.1039/c5cp06224a.

(99) Lin, P.; Yang, W.; Pedersen, L. C.; Negishi, M.; Pedersen, L. G. Searching for the minimum energy path in the sulfuryl transfer reaction catalyzed by human estrogen sulfotransferase: Role of enzyme dynamics. International Journal of Quantum Chemistry 2006, 106 (14), 2981–2998. DOI: 10.1002/qua.21123.

(100) Sousa, S. F.; Ribeiro, A. J. M.; Neves, R. P. P.; Brás, N. F.; Cerqueira, N. M. F. S. A.; Fernandes, P. A.; Ramos, M. J. Application of quantum mechanics/molecular mechanics methods in the study of enzymatic reaction mechanisms. WIREs Computational Molecular Science 2016, 7 (2). DOI: 10.1002/wcms.1281.

