## Supporting Information for "Mechanistic Basis of Sulfo Transfer in a PAPS-Independent ArylSulfate SulfoTransferases Revealed by Crystallography and QM/MM Simulations"

**Table of Contents**

S1. Abbreviations

S2. Crystal structure of *Desulfitobacterium hafniense*

S2.1. Dh_ASST_6 comparison with Ec_ASST_1

S3. Computational Details

S3.1. Structural and Bioinformatic analysis Loop modelling

S3.2. Protein Loop modelling

S3.3. Tunnel Analysis

S3.4. Constant pH Molecular Dynamics simulation

S3.5. Structure-guided generation of the reactant complex

S3.6. Molecular Dynamics simulation analysis

S4. Evaluation of the Steered QM/MM MD Protocol for the Second Catalytic Step

S5. Near attack conformation for the second catalytic step

### S1. Abbreviations

| | |
|---|---|
| **AMPSO** | 3-[(1,1-Dimethyl-2-hydroxyethyl)amino]-2-hydroxypropanesulfonic acid |
| **ASST** | Arylsulfate sulfotransferase |
| **ASTB** | Arylsulfate sulfotransferase B (from *Desulfitobacterium hafniense*) |
| **CpHMD** | Constant pH molecular dynamics |
| **CV** | Collective variable |
| **CV1** | Reaction coordinate for N–H bond formation (proton transfer) |
| **CV2** | Reaction coordinate for O–S bond formation (sulfo transfer) |
| **DFTB3** | Density Functional Tight Binding, third order |
| **GAFF** | General Amber Force Field |
| **HEPES** | 4-(2-Hydroxyethyl)-1-piperazineethanesulfonic acid |
| **MD** | Molecular dynamics |
| **MES** | 2-(N-Morpholino)ethanesulfonic acid |
| **MSA** | Multiple sequence alignment |
| **PAPS** | 3′-Phosphoadenosine-5′-phosphosulfate |
| ***p*NP** | *para*-Nitrophenol |
| ***p*NPS** | *para*-Nitrophenyl sulfate |
| **QM/MM** | Quantum mechanics/molecular mechanics |
| **RESP** | Restrained electrostatic potential |
| **RMSD** | Root-mean-square deviation |
| **UET** | Universal Evolutionary Trace |
| **LCO** | linear-combination-of-coordinates |

### S2. Crystal structure of Dh_ASST_6

**Table S1. Data collection and refinement statistics**

*Values in parentheses are for the highest-resolution shell.

| **Dataset name** | **DH_ASST_6** | **DH_ASST_6_*p*NPS** |
|---|---|---|
| Data collection | PX1 (07/09/2023) | PX1 (24/05/2024) |
| **Spage group** | $P2_12_12_1$ | $P2_12_12_1$ |

| | | |
|---|---|---|
| **Unit cell parameters** | a=100.67 b=113.23 c=151.22 | a=106.29 b=106.86 c=165.91 |
| **Resolution (Å)** | 49.20-1.9 (2.0-1.9) | 49.12-2.79 (2.87-2.79) |
| ***$R_{pim}$*** | 0.058 (1.12) | 0.100 (0.719) |
| **$CC_{1/2}$** | 0.994 (0.501) | 0.990 (0.391) |
| ***I/σ(I)*** | 11.6 (0.8) | 7.7 (0.9) |
| **Completeness (%)** | 99.8 (99.0) | 99.9 (98.9) |
| **Redundancy** | 13.9 (13.7) | 11.4 (10.9) |
| **No. Unique reflections** | 12,9279 (8526) | 47,512 (3426) |
| Refinement | | |
| **Resolution (Å)** | 49.20-1.9 | 49.12-2.79 |
| **No. atoms** | | |
| **Protein Chain 1** | 4949 | 4950 |
| **Protein Chain 2** | 4943 | 4958 |
| **Ligand/ion** | 5 | - |
| **Water** | 455 | 59 |
| ***Rwork/Rfree*** | 21.51/24.87 | 19.05/25.68 |
| **B-factors** | | |
| **Protein (Å²) Chain 1** | 38.37 | 62.40 |
| **Protein (Å²) Chain 2** | 38.98 | 65.39 |
| **Ligand/ion (Å²)** | 27.69 | |
| **Water (Å²)** | 37.15 | 44.20 |
| **R.m.s. deviations** | | |
| **Bond lengths (Å)** | 0.009 | 0.006 |
| **Bond angles (°)** | 1.518 | 1.477 |
| **Ramachandran plot** | | |
| **Favored (%)** | 97.03 | 94.20 |
| **Allowed (%)** | 2.47 | 4.97 |
| **Outliers (%)** | 0.50 | 0.83 |
| **PDB code** | **30FK** | **30FV** |

**Table S2. Structural alignment metrics of the catalytic and accessory domains of *Desulfitobacterium hafniense***

| | RMSD | Z score | Aligned residues |
|---|---|---|---|
| DH_ASST_6 catalytic vs EC_ASST_1 catalytic domain | 2.2[COOT] | 36.5 | 339/396 |
| DH_ASST_6 N-terminal vs EC_ASST_1 N-terminal | 2.13 [COOT] | 4.8 | 78/117 |
| DH_ASST_6 C-terminal vs EC_ASST_1 N-terminal | 2.13 [COOT] | 4.8 | 78/108 |
| DH_ASST_6 N-terminal vs | | | |
| 2qbw | 2.2[DALI] | 7.9 | 79/189 |
| 4lsd | 2.3 [DALI] | 7.2 | 81/98 |

#### S2.1. Dh_ASST_6 comparison with Ec_ASST_1

Dh_ASST_6 was compared with Ec_ASST_1 using the native and covalent-intermediate structures reported for the latter enzyme (PDB 3ELQ and 3ETT, respectively). The conserved histidine arrangement supports a shared sulfo-transfer core, whereas the surrounding pocket differs (**Figure S1**). In Ec_ASST_1, Tyr559, Asn358, Arg374, Thr501, and Tyr208 contribute to the sulfo-group environment; the corresponding region in Dh_ASST_6 includes Tyr487, Asn285, Arg301, Ile417, and Tyr194. The covalent Ec_ASST_1 intermediate in 3ETT contains the desulfated donor product 4-nitrophenol, whereas no corresponding ligand density was observed in Dh_ASST_6_*p*NPS. Differences involving Phe171, Ile500, Thr501, and Tyr208 in Ec_ASST_1 and the Tyr194/Tyr416/Ile417 region in Dh_ASST_6 could alter aromatic packing and ligand orientation. A spatial correspondence between residues does not necessarily imply an identical side-chain orientation or function. The ligand-occupancy difference alone does not establish a difference in affinity or release rate.

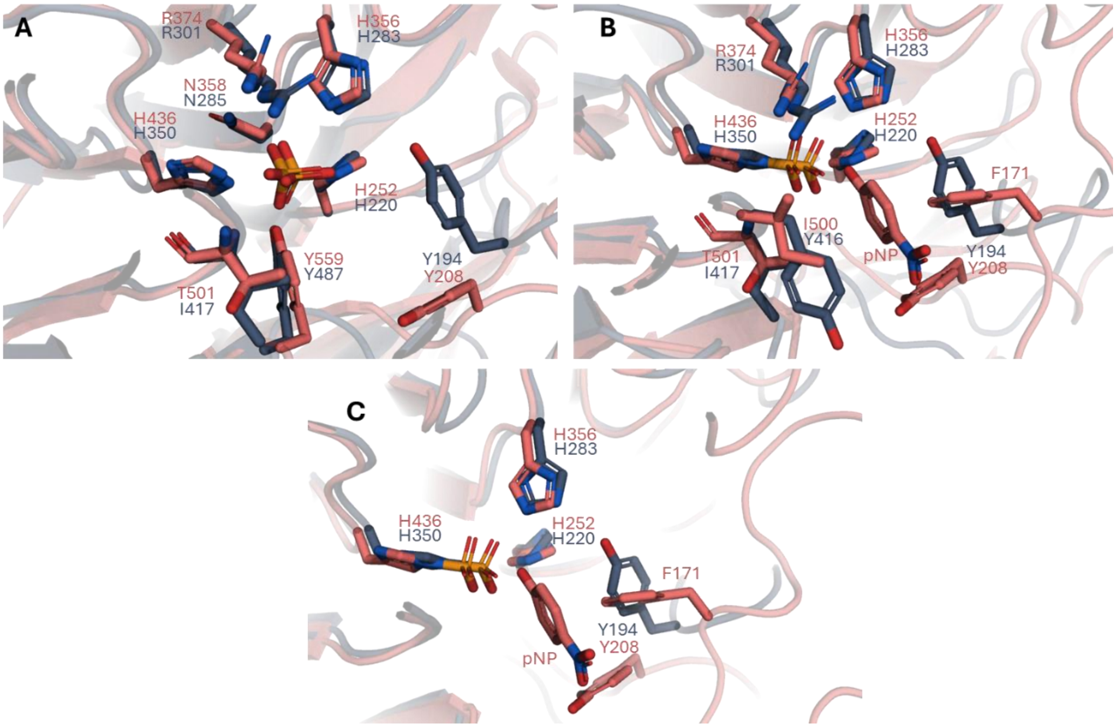

**Figure S1. Structural comparison of the active-site architecture of Dh_ASST_6 and Ec_ASST_1. (A)** Superposition of the active sites of Dh_ASST_6 (blue gray) and Ec_ASST_1 (salmon) showing the conserved sulfate-binding core centered around the catalytic histidine triad and neighbouring sulfate-coordinating residues. **(B)** Superposition of Dh_ASST_6_*p*NPS and Ec_ASST_1_*p*NPS highlighting differences in the donor-product-binding region. In both enzymes, the catalytic histidine is sulfated at Nε2, whereas the desulfated donor product *p*NP is resolved only in Ec_ASST_1. The comparison further reveals differences in the surrounding aromatic pocket, including Phe171 and Tyr208 in Ec_ASST_1 and the corresponding Tyr194/Tyr416 region in Dh_ASST_6. **(C)** Comparison of flexible loop regions near the active-site entrance, illustrating differences in loop placement and local pocket architecture between Dh_ASST_6 and Ec_ASST_1.

Residues 576–590 were unresolved in both Dh_ASST_6 crystal structures; residues 575 and 591 are the flanking anchors included in some loop displays. The entrance region differs from the corresponding flexible region in Ec_ASST_1 (**Figure S1C**). Such differences could affect substrate access, orientation, hydration, or product release, but the static structures do not establish a dynamic gating mechanism or rank the flexibility of the two enzymes.

### S3. Computational Details

#### S3.1. Structural and Bioinformatic analysis

To complement the structural characterization of Dh_ASST_6, sequence-based conservation analyses were performed to evaluate the evolutionary conservation of residues associated with the proposed active site. The 100 highest-scoring homologues identified by BLASTP searches against the non-redundant protein sequence database were included in the analysis. The selected sequences covered a broad identity range of 46.8–100%, and 98 of the 100 sequences covered at least 95% of the 626-residue Dh_ASST_6 sequence. A multiple-sequence alignment of Dh_ASST_6 homologues revealed that Tyr194, His220, His283, Asn285, Arg301, and His350 were invariant across all 100 analysed sequences. His220, His283, and His350 are therefore embedded within strongly conserved local sequence regions, supporting their proposed roles in the conserved catalytic architecture of bacterial arylsulfate sulfotransferases. Tyr416 and Ile417 were retained in all 99 sequences containing the corresponding region; the apparent conservation value of 99% results from one partial sequence that terminates before these positions. Tyr487 was present as tyrosine in 98 sequences, replaced by phenylalanine in one sequence, and absent from the same partial sequence. Thus, Tyr487 is strongly, but not absolutely, conserved (**Figure S2**). Conserved-domain analysis identified an N-terminal arylsulfotransferase Ig-like domain spanning residues 37–116 and the principal arylsulfotransferase domain spanning residues 127–488. All residues examined in the active-site conservation analysis are located within the latter domain, with Tyr487 positioned near its C-terminal boundary. This conservation pattern supports the functional relevance of these residues and is consistent with their proposed roles in catalysis and substrate binding. In particular, the conservation of the three histidine residues further supports the presence of a conserved histidine-centered catalytic architecture in this enzyme family.

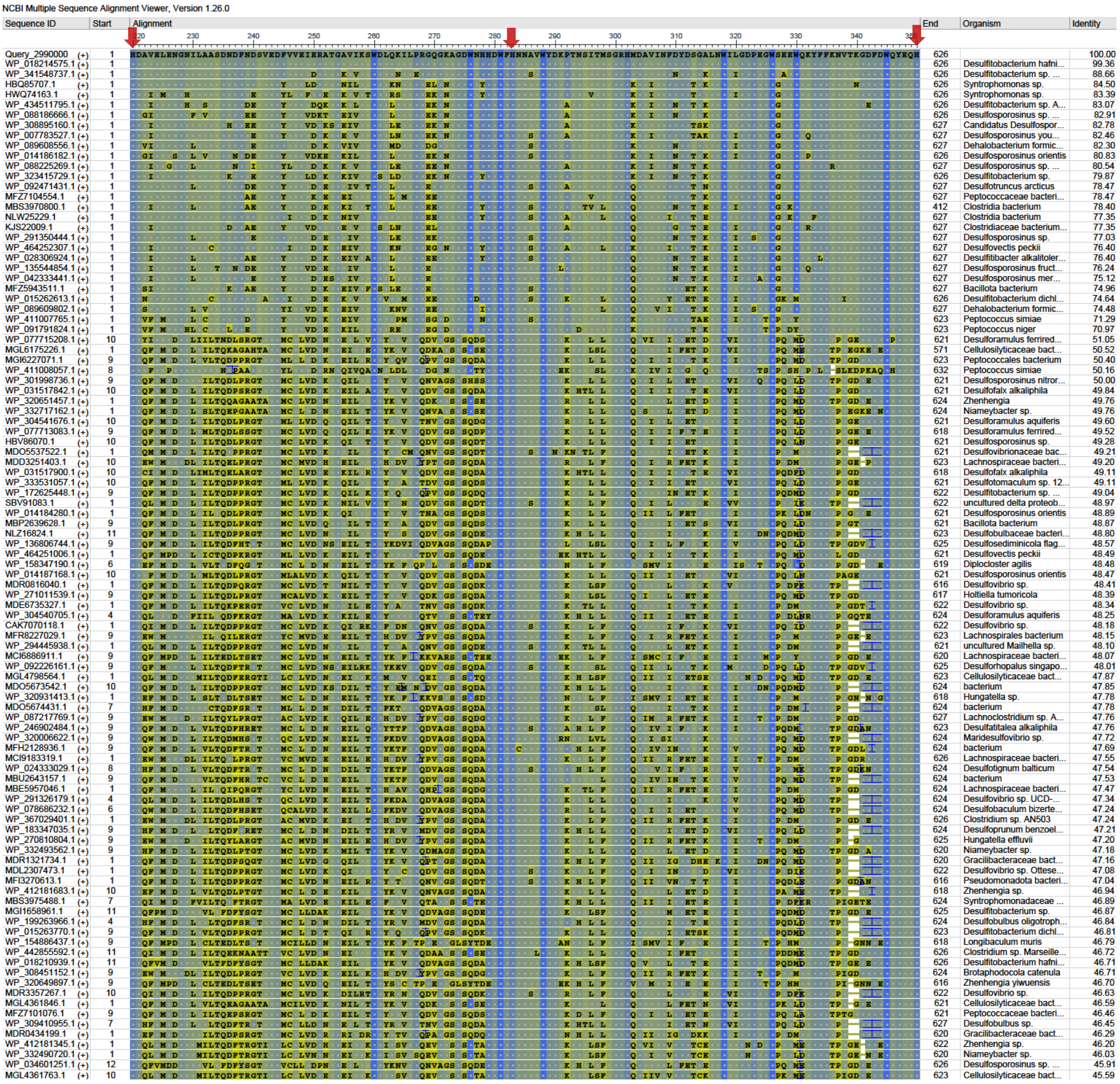

**Figure S2. Multiple-sequence alignment of Dh_ASST_6 homologues with evolutionary conservation analysis of key active-site residues**. Red arrows show the local sequence environments of His220, His283, and His350 together with residue conservation patterns derived from the alignment.

To examine conservation in structural context, evolutionary-importance scores from UET[4] were mapped onto the reconstructed Dh_ASST_6 structure, including the modelled loop. Highly ranked residues cluster in and around the internal catalytic cavity (**Figure S3**). The surface view locates this region within the more variable exterior, and the cartoon shows that distant sequence segments converge around the same catalytic center. The separate 100-sequence alignment identifies Tyr194, His220, His283, Asn285, Arg301, and His350 as invariant. Tyr416 and Ile417 are retained in all sequences covering those positions, whereas

Tyr487 is strongly but not absolutely conserved. **Table S3** distinguishes these sequence counts from their structural sulfate contacts.

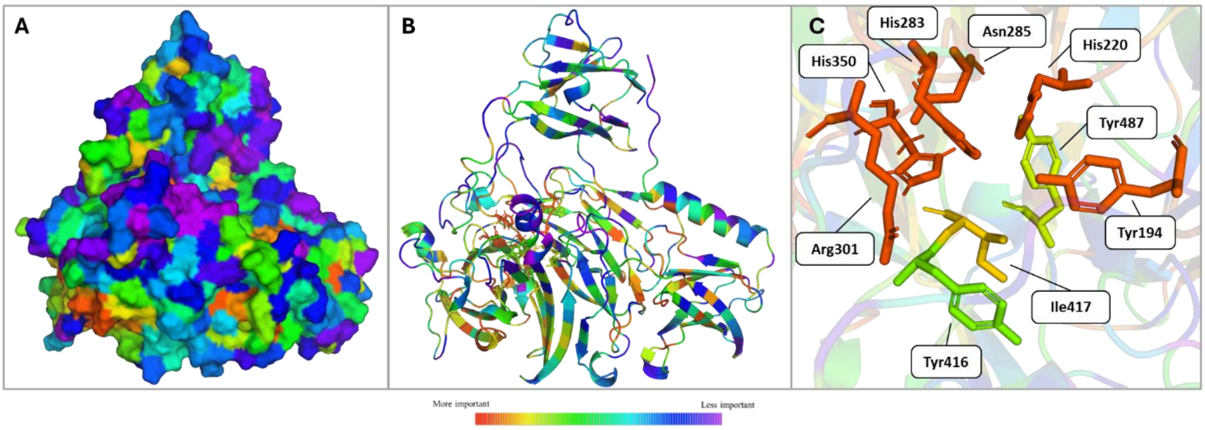

**Figure S3. Evolutionary trace analysis of Dh_ASST_6 based on the loop-modelled PDB structure. (A)** Surface representation of the full enzyme coloured according to evolutionary importance/conservation. The colour scale ranges from red/orange for highly important or conserved residues to blue/purple for less conserved residues. **(B)** Cartoon representation of the same structure, showing the spatial distribution of conserved residues throughout the enzyme fold. **(C)** Close-up view of the active-site region, highlighting highly conserved residues located within or near the catalytic pocket. The most conserved residues include Tyr194, His220, His283, Asn285, Arg301, and His350, while Tyr416, Ile417, and Tyr487 also show high evolutionary importance and are positioned close to the active-site environment.

**Table S3. Sequence conservation and structural context of residues surrounding the Dh_ASST_6 active site**. Conservation was calculated from the 100-sequence multiple-sequence alignment, with gaps included in the denominator. Sulfate-contact distances represent the mean of measurements from chains A and B of the crystallographic structure. All listed residues are located within the principal arylsulfotransferase domain spanning residues 127–488.

| **Residue** | **Occurrence in 100-sequence alignment** | **Conservation** | **Mean sulfate contact** | **Structural interpretation** |
|---|---|---|---|---|

| | | | | |
|---|---|---|---|---|
| **Tyr194** | 100 Tyr | 100% | Tyr OH···O, 3.92 Å | Lines the sulfate-binding pocket and contributes to its polar and aromatic environment; no direct sulfate hydrogen bond is observed |
| **His220** | 100 His | 100% | His NE2···O, 2.80 Å | Forms direct polar contact with sulfate and contributes to the conserved histidine-centered interaction network |
| **His283** | 100 His | 100% | His NE2···O, 2.86 Å | Directly contacts sulfate and is positioned to participate in acceptor activation during the second half-reaction |
| **Asn285** | 100 Asn | 100% | Asn ND2···O, 3.05 Å | Stabilizes the sulfate-binding region and the local hydrogen-bonding environment surrounding His283 |
| **Arg301** | 100 Arg | 100% | Arg NE···O, 3.94 Å | Contributes to the electrostatic and structural organization of the pocket, although no short direct contact is present in this conformation |
| **His350** | 100 His | 100% | His NE2···O, 2.61 Å | Shows the shortest polar contact to sulfate and corresponds to the sulfo-accepting catalytic histidine |
| **Tyr416** | 99 Tyr, 1 gap | 99% overall; 100% of non-gap sequences | Tyr OH···O, 7.74 Å | Forms part of the pocket wall and contributes mainly to pocket geometry rather than direct sulfate coordination |

| | | | | |
|---|---|---|---|---|
| **Ile417** | 99 Ile, 1 gap | 99% overall; 100% of non-gap sequences | Backbone N···O, 3.08 Å | Its backbone can interact with sulfate, whereas its side chain contributes to hydrophobic packing and pocket shape |
| **Tyr487** | 98 Tyr, 1 Phe, 1 gap | 98% overall; 99% of non-gap sequences | Tyr OH···O, 4.61 Å; aromatic C···O, 3.55 Å | Aromatic residue lining the pocket; the conservative Tyr-to-Phe substitution indicates that aromatic character may be more strongly constrained than the hydroxyl group |

The combined sequence and structural analyses therefore identify a strongly conserved active-site network centered on His220, His283, and His350, with additional invariant or near-invariant residues contributing to sulfate coordination and organization of the catalytic pocket.

**S3.2 Protein Loop modelling**

To reconstruct the unresolved region in the crystal structure, the missing loop comprising residues 576–590 (sequence VHDVTAACVSFEEQN) was modelled and sampled. Loop sampling in YASARA[5] generated an ensemble of candidate conformations that were subsequently subjected to local energy minimization. During minimization, the protein core was kept fixed, whereas the modelled loop and its adjacent anchor residues were allowed to relax. Ten independent modelling sets containing 31 minimized structures each were generated, resulting in a total ensemble of 310 loop-containing models.

To organize this conformational ensemble and identify recurrent loop geometries, pairwise RMSDs were calculated over residues 2–626 and subjected to hierarchical clustering. Because the protein core remained fixed during local refinement, the observed structural differences predominantly reflected variation in the reconstructed loop region. The pairwise RMSD matrix showed distinct block-like regions, indicating the presence of multiple related conformational families within the ensemble (**Figure S4A**). Clustering divided the 310 structures into 10 groups with markedly different populations.

Cluster 1 was the most populated and contained 72 structures, corresponding to 23.2% of the ensemble. Clusters 2–5 contained 47, 44, 40, and 39 structures, respectively, corresponding to 15.2%, 14.2%, 12.9%, and 12.6% of the ensemble. Together, the five largest clusters accounted for 242 structures, or 78.1% of the complete ensemble. The remaining clusters contained between 3 and 25 structures each. These results indicate that, despite the substantial variability of the reconstructed segment, most of the minimized models were concentrated within a limited number of recurrent conformational families. Cluster populations, maximum intracluster RMSDs, and representative structures are summarized in **Table S4**.

**Table S4. Summary of the RMSD-based clustering of the 310 energy-minimized Dh_ASST_6 loop models.** Cluster populations were calculated from the complete ensemble of 310 structures. The reported maximum RMSD values were calculated over residues 2–626 in the current analysis. Representative structures correspond to the medoid-like member having the lowest summed pairwise RMSD within each cluster.

| **Cluster** | **Structures, *n*** | **Population (%)** | **Maximum pairwise RMSD (Å)** | **Representative subjected to MD** |
|---|---|---|---|---|
| **1** | 72 | 23.2 | 1.291 | Yes |
| **2** | 47 | 15.2 | 1.295 | Yes |
| **3** | 44 | 14.2 | 1.432 | Yes |
| **4** | 40 | 12.9 | 1.531 | Yes |
| **5** | 39 | 12.6 | 1.654 | No |
| **6** | 25 | 8.1 | 1.683 | No |
| **7** | 16 | 5.2 | 1.493 | No |
| **8** | 14 | 4.5 | 1.583 | Yes |
| **9** | 10 | 3.2 | 1.519 | Yes |
| **10** | 3 | 1.0 | 1.270 | No |

A representative structure was identified for each cluster as the structure having the lowest summed pairwise RMSD to the other members of that cluster. Pairwise comparison of the ten representatives confirmed that the clustering procedure retained structurally distinct conformations for subsequent evaluation (**Figure S4B**).

Representatives from Clusters 1, 2, 3, 4, 8, and 9 were subsequently subjected to 200 ns classical MD simulations. Loop stability was evaluated from the RMSD of residues 575–591, comprising the modelled residues and one flanking anchor residue on each side, relative to the corresponding minimized starting structure. The trajectories showed distinct degrees of structural relaxation and conformational stability (**Figure S4C**). The Cluster 1 representative underwent limited initial relaxation and subsequently maintained the lowest and most stable RMSD plateau, predominantly around 2 Å. The Cluster 4 representative also retained comparatively low RMSD values during the later part of the trajectory. Clusters 2 and 3 underwent larger intermediate rearrangements, whereas the Cluster 8 representative showed pronounced fluctuations. Cluster 9 displayed the greatest conformational drift, with RMSD values frequently exceeding 6 Å and reaching approximately 9–10 Å during parts of the trajectory.

The Cluster 1 representative was additionally inspected within the crystallographic model with respect to connectivity between the resolved anchor residues, steric compatibility with the surrounding protein environment, and consistency with the available electron density near the loop boundaries. The selected conformation formed a continuous and sterically plausible connection between the resolved ends of the missing segment and remained within the spatial region compatible with the experimental map. Because residues 576–590 were unresolved in the crystal structure, the electron density does not define a unique conformation for the complete loop. Nevertheless, the combination of its high cluster population, comparatively low conformational drift during MD, and compatibility with the crystallographic environment supported selection of the Cluster 1 representative as the starting model for the subsequent computational analyses.

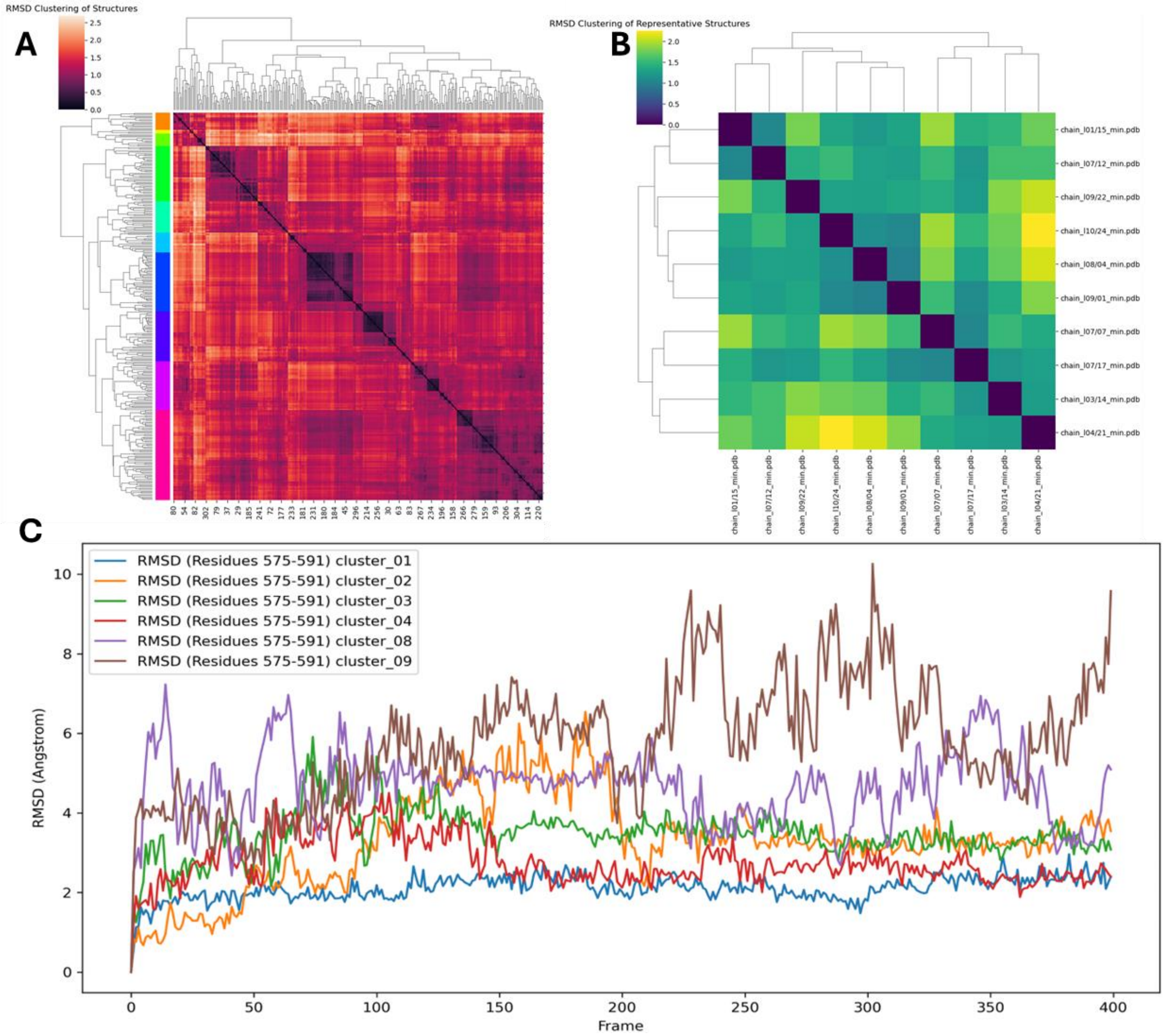

**Figure S4: RMSD-based clustering and MD evaluation of the modelled Dh_ASST_6 loop conformations**. **(A)** Hierarchically clustered pairwise RMSD matrix of the 310 energy-minimized loop-containing models. In the current analysis, RMSDs were calculated over residues 2–626. The coloured annotation denotes assignment to the 10 resulting clusters. **(B)** Pairwise RMSD matrix of the representative structures selected from the 10 clusters. **(C)** RMSD profiles of residues 575–591, comprising the modelled loop and its flanking anchor residues, during 200 ns MD simulations of representatives from Clusters 1, 2, 3, 4, 8, and 9. RMSD values were calculated relative to the corresponding minimized starting structure.

The reconstructed loop is positioned at the entrance of the catalytic cavity, spatially separated from the conserved histidine-centered catalytic core. Its pronounced conformational heterogeneity during modelling and MD suggests that it may function as a flexible boundary of the active-site pocket. By sampling more open and more closed conformations, the loop could modulate substrate access, solvent exposure, and product release without directly participating in sulfuryl transfer. The selected Cluster 1 conformation therefore represents a plausible starting geometry for the simulations, while the observed mobility indicates a potential gating role that remains to be experimentally established.

**S3.3 Tunnel Analysis**

Tunnel analysis of the loop-modelled Dh_ASST_6 structure was performed using the CAVER 3.0 plugin in PyMOL[7, 8] The search was initiated from coordinates x=14.383, y=5.705, and z=8.637 Å within the ligand-binding region, using a probe radius of 0.7 Å, a shell radius of 3.0 Å, and a shell depth of 4.0 Å. After removal of 128 redundant pathways, 29 tunnels were retained and grouped into eight clusters by average-link hierarchical clustering using a threshold of 3.5 Å. The lowest-cost representative from each cluster was retained for subsequent analysis, and the resulting cluster populations and geometric descriptors are summarized in **Table S5**.

**Table S5. Quantitative characteristics of the eight tunnel clusters identified in the loop-modelled Dh_ASST_6 structure.** The cluster population refers to the number of the 29 nonredundant pathways assigned to each cluster. The lowest-cost tunnel within each cluster was retained as the representative.

| CAVER cluster | Number of tunnels | Population (%) | Bottleneck radius (Å) | Length (Å) | Curvature |
|---|---|---|---|---|---|
| **1** | 2 | 6.9 | 0.746 | 23.90 | 1.355 |
| **2** | 2 | 6.9 | 0.746 | 28.14 | 1.837 |
| **3** | 8 | 27.6 | 0.746 | 35.54 | 1.433 |
| **4** | 12 | 41.4 | 0.746 | 27.30 | 1.413 |
| **5** | 1 | 3.4 | 0.704 | 39.69 | 1.775 |
| **6** | 2 | 6.9 | 0.741 | 62.33 | 1.617 |

| | | | | | |
|---|---|---|---|---|---|
| **7** | 1 | 3.4 | 0.704 | 51.98 | 1.933 |
| **8** | 1 | 3.4 | 0.713 | 62.85 | 1.413 |

Clusters 3 and 4 accounted for 20 of the 29 pathways, corresponding to 69.0% of the detected ensemble, whereas the remaining clusters contained only one or two tunnels each. The representative pathways varied substantially in length, from 23.9 to 62.9 Å, but displayed similarly narrow bottleneck radii of 0.70–0.75 Å. Because these values are close to the applied probe radius, the detected routes should be interpreted as geometrically accessible pathways within the analysed structure rather than as evidence that complete donor or acceptor substrates can pass through each tunnel without conformational rearrangement. Four structurally distinct pathways were selected for visualization and comparison in **Figure S5**.

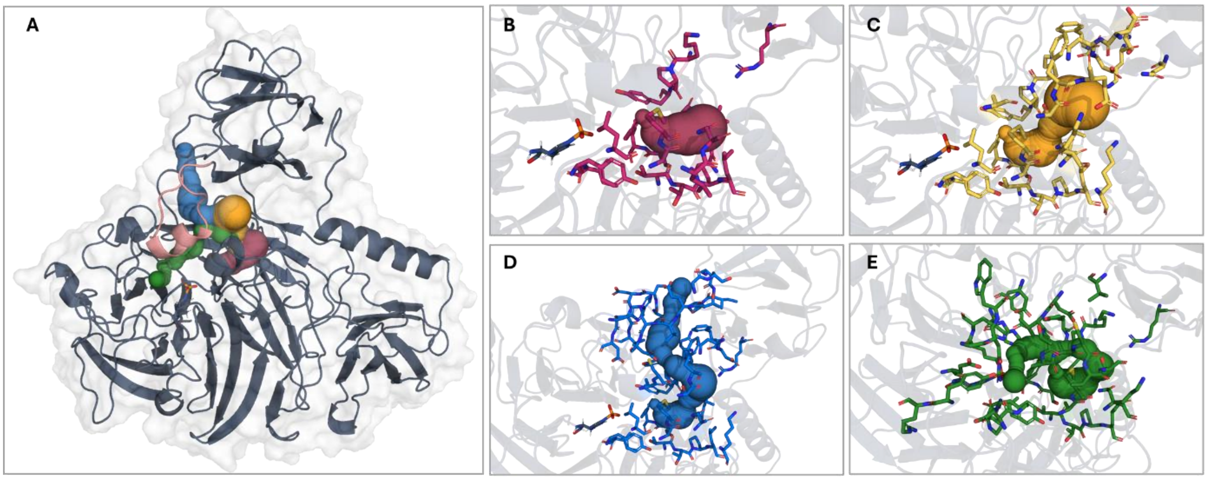

**Figure S5: Tunnel analysis of Dh_ASST_6. (A)** Overview of four representative pathways selected from the eight tunnel clusters identified in the complete analysis. The protein is shown as a dark-gray cartoon with its molecular surface in white. The pathways are represented by coloured spheres, and the modelled loop comprising residues 576–590 is highlighted in pink. **(B–E)** Individual representations of displayed pathways A–D together with their tunnel-lining residues. Quantitative properties of all eight CAVER clusters are reported in **Table S5**, and the lining residues of the four displayed pathways are listed in **Table S6**.

Their complete lining-residue sets and contributions from the reconstructed loop are provided in **Table S6**. The pathways share an inner amphipathic region formed primarily by Thr144–Leu147, Glu170–Trp172, Pro193, Tyr194, Tyr416, Ile417, Leu446, Ala448, Met449, Asn452, and Arg548, but diverge toward different surface openings.

**Table S6. Residue composition and structural characteristics of the four pathways displayed in Figure S5.** Residue 627 was used to define or mark the ligand-binding region and is therefore not included among the protein tunnel-lining residues. Superscript "bb" denotes residues contributing primarily through backbone atoms.

| Displayed pathway | Tunnel-lining residues | Modelled-loop residues, 576–590 | Notable active-site residues | Structural interpretation |
|---|---|---|---|---|
| **A** | Thr144, Thr145, Pro146, Leu147, Asn168, Leu169, Glu170, Ser171, Trp172, Lys192, Pro193, Tyr194, Tyr416, Ile417, Leu446, Ala448, Met449, Leu451, Asn452, Arg548 | None | Tyr194, Tyr416, Ile417 | Compact amphipathic pathway that is independent of the reconstructed loop |
| **B** | Thr144, Thr145, Pro146, Leu147, Leu169, Glu170, Ser171, Trp172, Lys192, Pro193, Tyr194, Tyr416, Ile417, Leu446, Ala448, Met449, Asn452, Lys453, Lys455, Asn535, Arg548, Ser550, Ala582, Ser585, Phe586, Glu587, Asp591, Asn592, Asp593, Phe594 | Ala582^bb, Ser585, Phe586^bb, Glu587 | Tyr194, Tyr416, Ile417 | Related amphipathic pathway with a more polar outer region and limited contribution from the distal loop |
| **C** | Thr144, Thr145, Pro146, Leu147, Leu169, Glu170, | Val576, His577, Asp578, Ala582, | His279, Asp280, | Strongly amphipathic |

| | | | | |
|---|---|---|---|---|
| | Ser171, Trp172, Gln191, Lys192, Pro193, Tyr194, Tyr195, Phe238, Asn239, Asp240, Ser241, Val242, His279, Asp280, Tyr416, Ile417, Leu446, Ala448, Met449, Asn452, Lys455, Arg548, Gly554, Ser555, Asn571, Ile572, Pro573, Thr574, Glu575, Val576, His577, Asp578, Ala582, Cys583, Ser585, Phe586, Asn592, Asp593, Phe594 | Cys583, Ser585, Phe586 | Tyr194, Tyr416, Ile417 | alternative pathway with substantial contribution from the reconstructed loop |
| **D** | Thr144, Thr145, Pro146, Leu147, Leu169, Glu170, Ser171, Trp172, Lys192, Pro193, Tyr194, Tyr195, Glu243, Gly274, Asp275, Trp276, Asp280, His283, Arg301, His302, Glu348, sulfonated His350, Thr367, Tyr368, Pro415, Tyr416, Ile417, Asp444, Gly445, Leu446, Ala448, Met449, Asn452, Lys453, Arg548, Val576, Asp578, Val579, Thr580, Ala581, Ala582, Cys583, Val584, Ser585 | Val576, Asp578, Val579, Thr580^bb, Ala581, Ala582, Cys583^bb, Val584, Ser585 | His283, Arg301, His302, sulfonated His350 | Pathway passing closest to the catalytic center and showing the strongest continuous contribution from the reconstructed loop |

The reconstructed loop does not contribute to displayed pathway A, contributes only locally to pathway B, and forms increasingly extensive boundaries along pathways C and D. Pathway D additionally passes close to His283, Arg301, His302, and sulfonated His350 and therefore approaches the reactive center most directly. These results indicate that residues 576–590 do not form a universal bottleneck but may modulate the geometry, hydration, and flexibility of selected peripheral access routes.

#### S3.4 Constant pH Molecular Dynamics simulation

Constant-pH molecular dynamics simulations (CpHMD)[6, 9] were used to evaluate the protonation preferences of the catalytic histidines His220, His283, and His350 at pH 6.5. The three residues displayed distinct and strongly defined protonation-state populations (**Figure S6** and **Table S7**). His283 remained almost exclusively in the doubly protonated, positively charged HIP state. In contrast, His220 and His350 were predominantly neutral, but favoured different neutral tautomers. His220 adopted primarily the Nε-protonated HIE state, with a population of 0.9862, whereas His350 was almost exclusively Nδ-protonated in the HID state, with a population of 0.9998.

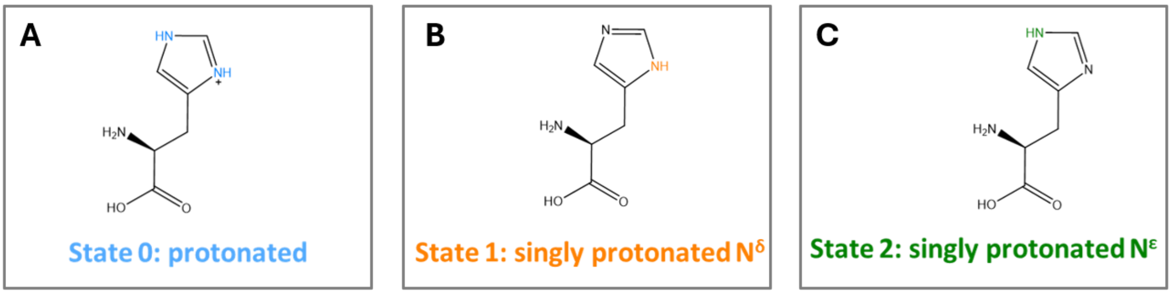

**Figure S6. Schematic representation of the histidine protonation states used in the constant-pH molecular dynamics analysis. (A)** State 0 corresponds to the doubly protonated, positively charged HIP form; **(B)** State 1 corresponds to the neutral Nδ-protonated HID tautomer; and **(C)** State 2 corresponds to the neutral Nε-protonated HIE tautomer.

**Table S7. Protonation-state populations and predicted pKa values of the catalytic histidines at pH 6.5.** HIP denotes the positively charged state protonated at both Nδ1 and Nε2. HID and HIE are neutral tautomers protonated at Nδ1 and Nε2, respectively.

| Residue | HIP population | HID population | HIE population | Dominant state | Protonated nitrogen in | Formal charge | Predicted pK(_a) |
|---|---|---|---|---|---|---|---|

| | | | | | dominant state | | |
|---|---|---|---|---|---|---|---|
| **His220** | 0.001235 | 0.012530 | 0.986235 | HIE | Nε2–H; Nδ1 unprotonated | 0 | 3.592 |
| **His283** | 1.000000 | 0.000000 | 0.000000 | HIP | Nδ1–H and Nε2–H | +1 | Not resolved; persistently protonated |
| **His350** | 0.000155 | 0.999845 | ≈0.000000 | HID | Nδ1–H; Nε2 unprotonated | 0 | 2.690 |

The predicted pKa values of His220 and His350 were 3.59 and 2.69, respectively, consistent with their predominantly neutral states at pH 6.5. His283 remained fully protonated throughout the sampled pH range, preventing determination of a finite pKa value under the applied conditions. The dominant microstate of the substrate-free enzyme therefore consists of HIE220, HIP283, and HID350.

This protonation pattern is compatible with the proposed organization of the catalytic site. The positively charged His283 may stabilize the anionic sulfate-binding environment in the substrate-free state, whereas the HID state of His350 leaves Nε available for formation of the covalent sulfo-histidine intermediate. The CpHMD results therefore provided the initial protonation-state assignment for subsequent structural modelling, while the protonation of His283 was adjusted where required to represent the reactive state of the second half-reaction.

#### S3.5 Structure-guided generation of the reactant complex

An initial reactant complex for the second half-reaction was generated by structure-guided placement of phenol into the active site of the sulfonated Dh_ASST_6 structure. Protonation states were assigned to represent a catalytically competent configuration for transfer of the sulfo group from His350 to the acceptor rather than the dominant microstate of the substrate-free enzyme obtained from constant-pH simulations. His283 was therefore modelled as the neutral Nδ1-protonated tautomer, leaving Nε2 available to accept the phenolic proton. His220 was retained in the neutral Nε2-protonated form, His350 was represented as the sulfonated catalytic intermediate, and phenol was modelled in its neutral state.

Because the Dh_ASST_6 crystal structure did not contain a bound acceptor, the initial phenol orientation was derived from a homologous ASST structure containing bound 4-nitrophenol. The Dh_ASST_6 structure was aligned with the reference enzyme in PyMOL[8, 10] by superposition of the conserved active-site residues corresponding to His220, His283, Arg301, and His350. Phenol was then fitted onto the shared phenolic ring and hydroxyl group of the reference 4-nitrophenol ligand, excluding the nitro substituent absent from phenol. This procedure oriented the phenolic hydroxyl group toward His283 and placed the acceptor oxygen in proximity to the sulfur center of sulfonated His350, consistent with the proposed proton-transfer and sulfuryl-transfer geometry.

The resulting complex was inspected for steric compatibility with the surrounding pocket and the sulfo-enzyme intermediate. The alignment-derived pose was used only as a mechanistically informed starting configuration and was subsequently relaxed and refined during classical MD simulations.

#### S3.6 Molecular Dynamics simulation analysis

The conformational stability of the Dh_ASST_6 sulfo-enzyme–phenol complex was assessed across six independent classical MD simulations[6, 11, 12]. Each trajectory was aligned to the minimized starting structure using protein backbone atoms to remove global translation and rotation, after which backbone RMSD values were calculated using MDAnalysis[13] and custom Python scripts. Following an initial relaxation, all replicas remained within approximately 1–2 Å of the reference structure for most of the simulations, indicating that the overall protein fold remained stable (**Figure S7**). Some replica-dependent deviations were observed at later simulation times, most notably in Runs 1 and 3, but these did not indicate global unfolding.

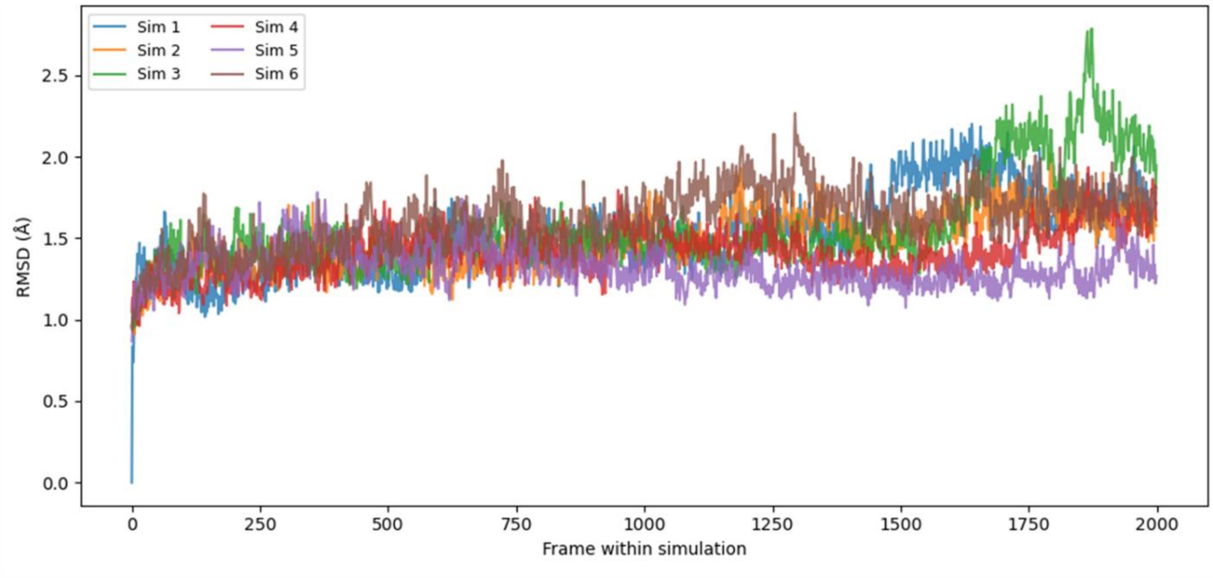

**Figure S7: Backbone RMSD profiles of six independent classical MD simulations of the** Dh_ASST_6 sulfo-enzyme–phenol complex. RMSD values were calculated for protein backbone atoms after alignment to the minimized starting structure. Runs 1–6 are shown separately**.**

To evaluate sampling of geometries relevant to the second half-reaction, two active-site distances were monitored throughout the concatenated trajectories: the distance between the phenolic oxygen and the sulfur atom of sulfonated His350, $d_{O-S}$, and the distance between the phenolic proton and the proton-accepting Nε2 atom of His283, $d_{H-N}$. These coordinates describe the simultaneous approach of the acceptor oxygen toward the sulfonyl sulfur and positioning of the phenolic proton for transfer to His283. A geometric filter of $d_{O-S} < 5.5$Å and $d_{H-N} < 2.2$Å was used to identify pre-reactive conformations. Only 3.91% of the analysed frames satisfied both criteria simultaneously (**Figure S8**), showing that this geometry constitutes a minor subpopulation of the unbiased classical MD ensemble. Most conformations satisfied neither condition or fulfilled only one of the two distance requirements, indicating that productive positioning requires correlated rearrangement of the acceptor relative to both His283 and the sulfo-enzyme intermediate.

The limited population of the filtered subensemble is consistent with substantial positional freedom of phenol within the active-site pocket. Because the initial acceptor pose was generated by structural alignment and phenol lacks the substituent present in the reference 4-nitrophenol ligand, it may experience fewer orientational constraints in the pocket. However, the present simulations do not establish that this chemical difference alone accounts for the observed conformational heterogeneity.

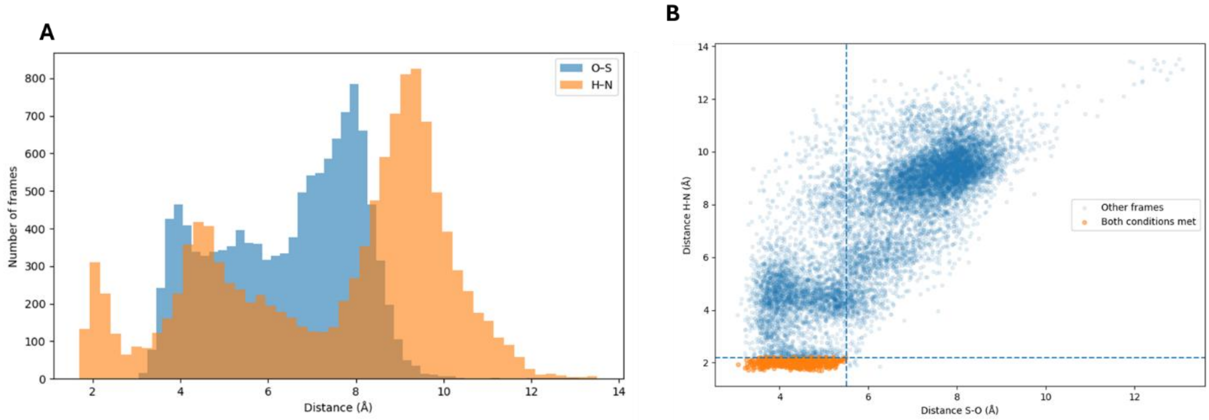

**Figure S8. Active-site distance distributions in the classical MD ensemble of the Dh_ASST_6 sulfo-enzyme–phenol complex. (A)** Distributions of the phenolic O···S distance to sulfonated His350 and the phenolic H···Nε2 distance to His283. **(B)** Two-dimensional distribution of the same distances over all analysed frames. Dashed lines indicate the thresholds used to define the pre-reactive subensemble, $d_{O-S} < 5.5$ Å and $d_{H-N} < 2.2$ Å. Frames satisfying both criteria are highlighted in orange and account for 3.91% of the total ensemble.

To characterize the broader conformational ensemble, the aligned backbone coordinates from the combined trajectories were subjected to hierarchical clustering. Five clusters were identified, and a representative structure was selected from each cluster as the sampled frame closest to the corresponding cluster center. The cluster-sorted all-versus-all RMSD matrix showed substantial internal similarity across the ensemble, with localized blocks corresponding to the identified conformational groups (**Figure S9A**). Pairwise RMSDs between the five representatives ranged from approximately 1.3 to 1.8 Å, indicating that the clusters describe moderate rearrangements within a conserved global fold rather than highly distinct protein conformations (**Figure S9B**).

Principal component analysis provided a complementary projection of the sampled conformational space. The first two components accounted for 13.8% and 12.6% of the positional variance, respectively. Cluster 5 occupied the broad central region of the projection, whereas Clusters 1 and 3 extended toward more separated regions of conformational space. Considerable overlap remained between several clusters, consistent with a continuous and partially interconverting ensemble rather than sharply separated structural states **(Figure S9C)**. All analyses were performed using MDAnalysis, NumPy, pandas, SciPy, scikit-learn, and Matplotlib with in-house Python scripts.

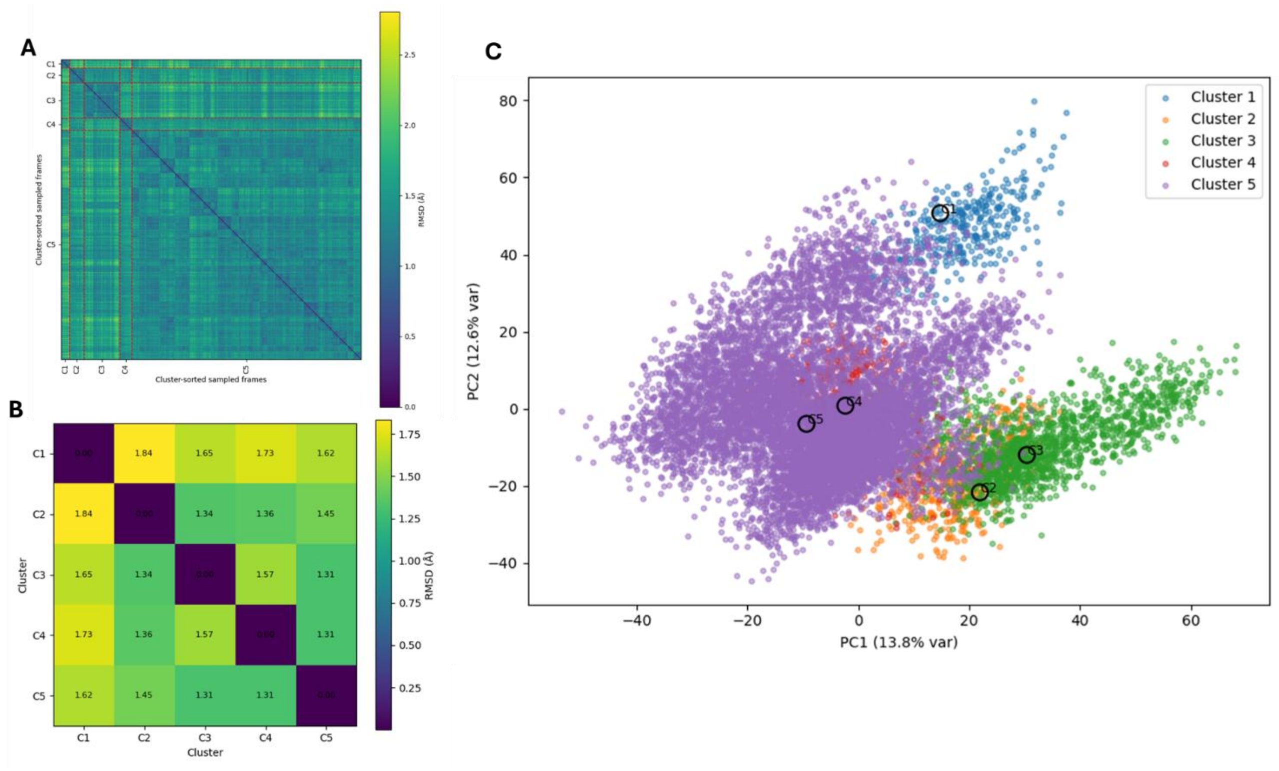

**Figure S9: Conformational clustering of the combined classical MD trajectories of the Dh_ASST_6 sulfo-enzyme–phenol complex. (A)** Cluster-sorted all-versus-all backbone RMSD matrix of the sampled frames. Cluster boundaries are indicated along the axes. **(B)** Pairwise backbone RMSD matrix of the five representative structures, with values reported in Å. **(C)** Projection of the trajectory frames onto the first two principal components, accounting for 13.8% and 12.6% of the positional variance, respectively. Frames are coloured according to cluster assignment, and the representative structure of each cluster is indicated by a black circle.

### S4. Evaluation of the Steered QM/MM MD Protocol for the Second Catalytic Step

To establish a suitable steered QM/MM MD[6, 11, 12] protocol for the second catalytic, rate limiting step, a series of preliminary calculations was performed in which both the QM region[14] and the collective-variable (CV) definition were systematically varied. The aim was to identify a setup that (i) provides a chemically meaningful description of the active-site environment, (ii) reaches the product region within the applied 2 ps steering window, and (iii) yields an interpretable Jarzynski-derived free-energy profile. Because unbiased classical MD sampling indicated that catalytically competent pre-reactive conformations are only rarely populated, starting structures for the steered QM/MM MD calculations were selected using a geometric filter based on the key distances associated with sulfo transfer and proton transfer. Specifically, frames were retained when the O–S distance between the acceptor oxygen atom and the sulfur atom of the sulfated His350 intermediate was below 5.5 Å and, simultaneously, the H–N distance between the transferring proton and the catalytic nitrogen atom of His283 was below 2.2 Å. These criteria identify conformations close to a near-attack arrangement suitable for initiation of the second half-reaction. In total, 314 trajectories were included in the final steered QM/MM MD analysis.

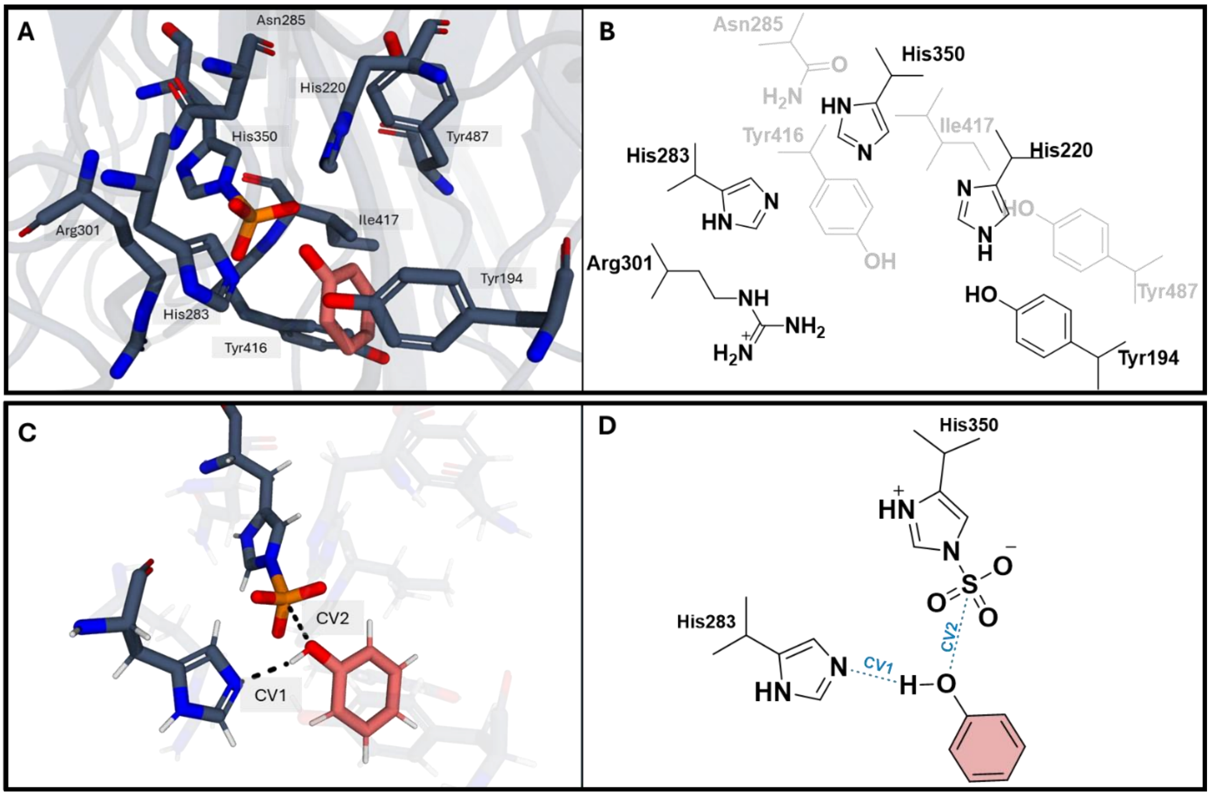

**Figure S10: Definition of the active-site QM region and collective variables used for the steered QM/MM MD simulations of the second catalytic step. (A)** Active-site view of the selected expanded QM region used in the final production setup. The acceptor ligand and catalytically relevant residues included in the QM region are shown as sticks, whereas hydrogens are omitted for clarity. **(B)** Schematic representation of the corresponding protonation states of the residues included in the QM region. **(C)** Three-dimensional representation of the two distance-based collective variables used in the steered QM/MM MD simulations. CV1 describes the N–H bond formation associated with proton transfer to His283, and CV2 describes the O–S bond formation between the acceptor oxygen atom and the sulfur atom of the sulfonated His350 intermediate. **(D)** Two-dimensional schematic of the same collective variables, highlighting the geometric definition of CV1 and CV2 used to drive the coupled proton-transfer and sulfo-transfer process.

Two QM region definitions were evaluated. The minimal QM region comprised the acceptor ligand together with His350 and His283 and had a total QM-region charge of 0. A second, expanded QM region additionally included Arg301, His220, Tyr487, Asn285, Tyr416, Ile417, and Tyr194, giving a total QM-region charge of +1. The expanded QM region was considered chemically more complete because it captures the immediate hydrogen-bonding and electrostatic environment around the reacting center and is therefore better suited to describe charge redistribution during the coupled proton-transfer and sulfo-transfer process.

Different CV definitions were then tested. In the first approach, the reaction was described by two distance-based CVs corresponding to N–H bond formation and O–S bond formation. In the second approach, a third distance-based CV describing S–N bond cleavage was added. In addition, linear-combination-of-coordinates (LCO) type CVs based on combined angular/geometric terms were examined. The tested setups and their qualitative outcomes are summarized in **Table S3**.

**Table S8: Tested steered QM/MM MD setups for the second catalytic step**.

| Setup | QM region | QM charge | CV definition | Approx. barrier (kcal/mol) | Outcome |
|---|---|---|---|---|---|
| **1** | Minimal QM region | 0 | 2 distance CVs: N–H formation, O–S formation | ~13.0 | Reaction completed; minimal QM description |
| **2** | Minimal QM region | 0 | 3 distance CVs: N–H formation, | ~16.9 | Reaction completed; |

| | | | | | |
|---|---|---|---|---|---|
| | | | O–S formation, S–N cleavage | | higher apparent barrier |
| **3** | Expanded QM region | +1 | 2 distance CVs: N–H formation, O–S formation | selected ~14.0 | Reaction completed; selected production setup |
| **4** | Expanded QM region | +1 | 3 distance CVs: N–H formation, O–S formation, S–N cleavage | ~23.0 | Reaction completed; strongly increased apparent barrier |
| **5** | Expanded QM region | +1 | 2 LCO CVs: O–H–N and N–S–O | ~15.3 (only preorganization) | Reaction not completed within 2 ps |
| **6** | Expanded QM region | +1 | 1 LCO CV: N–S–O | ~22.1 (only preorganization) | Reaction not completed within 2 ps |

Among the tested protocols, the expanded QM region combined with two distance-based CVs for N–H and O–S bond formation was selected for production calculations. Although the minimal QM region with the same CV definition produced a somewhat lower apparent barrier, it was considered less reliable because it neglects several catalytically relevant residues expected to stabilize the evolving charge distribution and substrate positioning during the reaction. By contrast, the expanded QM region includes the conserved active-site environment surrounding the catalytic center and therefore provides a more realistic description of the chemical event. Addition of a third CV, S–N bond cleavage, led to substantially higher barriers for both QM region definitions. The inclusion of the cleavage coordinate over constrains the reaction pathway. Likewise, the LCO-based CV definitions were less suitable under the present steering conditions, since these simulations did not reach a product-like state within the 2 ps simulation time and were therefore not pursued further. The selected production setup thus represented the best compromise between chemical completeness of the QM region, successful progression to the product region within the applied steering interval, and a continuous, mechanistically interpretable nonequilibrium pathway.

To assure the robustness of the selected protocol, convergence of the Jarzynski-derived free-energy[15] profile was evaluated by reconstructing the profile using increasing numbers of trajectories (**Figure S11 A+B**). Profiles generated from 50 and 100 trajectories yielded barriers of 10.92 and 10.93 kcal/mol, respectively, whereas inclusion of 200, 250, and all 314 trajectories gave values of 9.57, 9.57, and 9.58 kcal/mol. Thus, the reconstructed profile converges once approximately 200 trajectories are included.

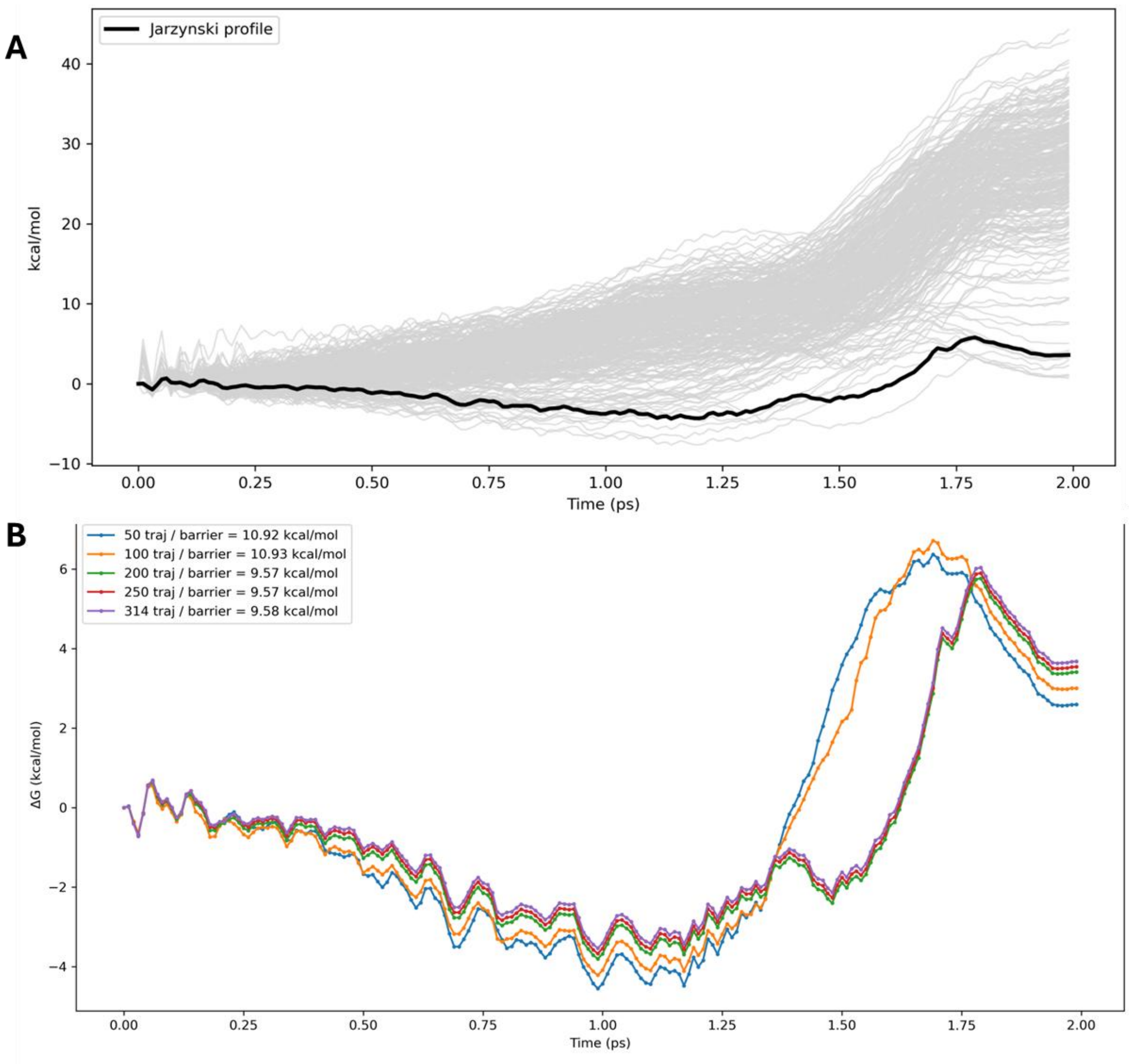

**Figure S11: Nonequilibrium work profiles and convergence of the Jarzynski-derived free-energy reconstruction for the selected steered QM/MM MD setup of the second catalytic step. (A)** Individual nonequilibrium work trajectories obtained from the steered

QM/MM MD simulations are shown in grey, and the corresponding Jarzynski-derived free-energy profile is shown as a black line. The selected production protocol employed the expanded QM region and two distance-based collective variables describing N–H bond formation and O–S bond formation. **(B)** Convergence of the Jarzynski free-energy profile as a function of the number of included trajectories. Profiles reconstructed from subsets of 50 (blue), 100 (orange), 200 (green), 250 (red), and the full trajectory (purple) set are overlaid.

### S5. Near attack conformation for the second catalytic step

To further characterize the early phase of the second catalytic step, the structural features of the shallow intermediate identified in the Jarzynski-derived free-energy profile were analyzed in detail. This intermediate, state III, is interpreted as a near-attack conformation that forms after active-site reorganization and immediately precedes the main bond-reorganization event. In this state, the acceptor has already adopted a reactive orientation toward the sulfur center of the sulfated His350 intermediate, while full transfer of the sulfo group has not yet occurred.

The three-dimensional structure of state III shows a preorganized arrangement in which the acceptor oxygen is positioned toward the sulfur center in a geometry compatible with productive sulfo transfer, while His283 is aligned to mediate proton transfer. The corresponding interatomic distances and angular parameters further support the assignment of this configuration as a catalytically competent pre-reactive state. Thus, state III represents the endpoint of the preorganization phase and the immediate for the dominant barrier-crossing event observed later along the reaction coordinate.

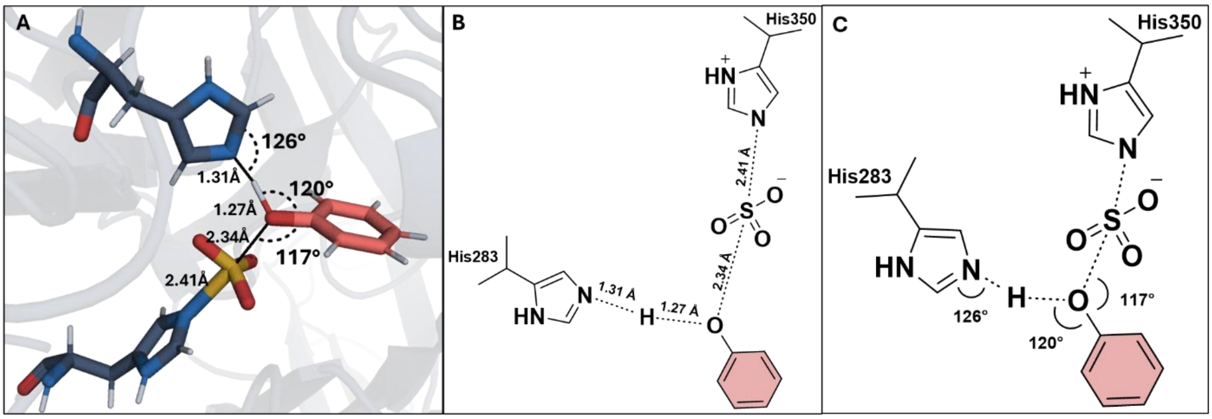

**Figure S12. Structural characterization of the near-attack conformation (state III) for the second catalytic step. (A)** Three-dimensional view of state III, showing the preorganized reactive arrangement formed after active-site preorganization and immediately preceding chemical transformation. Relevant distances and angles associated with the sulfo-transfer geometry are indicated. **(B)** Two-dimensional schematic of the same state highlighting the key interatomic distances used to describe the near-attack conformation. **(C)** Two-dimensional schematic highlighting the angular parameters that further define the reactive geometry of state III.

To examine how this structural reorganization is reflected in collective-variable space, all sampled configurations were projected onto the two-dimensional space defined by CV1 (H–N distance) and CV2 (O–S distance). The resulting occupancy map reveals a narrow and continuous pathway connecting reactant-like and product-like regions, demonstrating that the reaction proceeds along a well-defined configurational channel without evidence for competing routes. This observation is consistent with the free-energy profile reconstructed from the Jarzynski analysis.

To complement this structural representation, all sampled configurations were projected onto the two-dimensional space defined by CV1 (H–N distance) and CV2 (O–S distance). **Figure S13A** shows the occupancy distribution in this collective-variable space, whereas **Figures S13B** and **S13C** show the same projection coloured by simulation time and accumulated work, respectively.

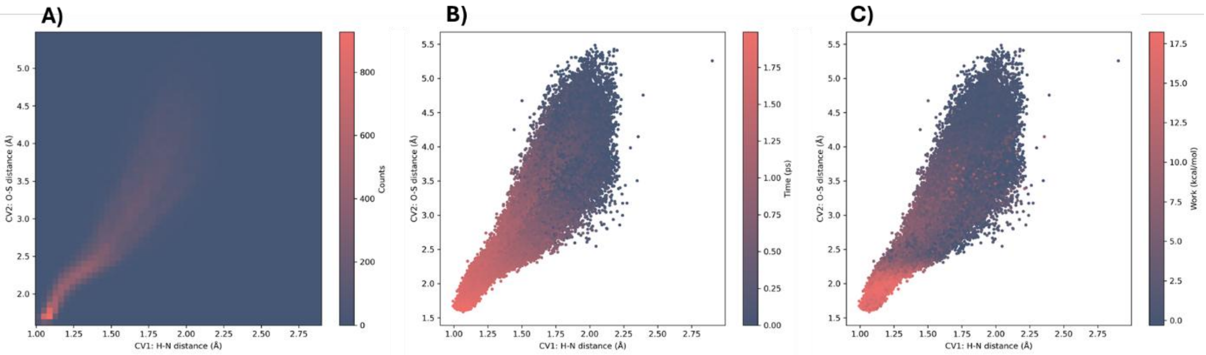

**Figure S13**. Distribution of sampled configurations in the two-dimensional collective variable (CV) space defined by CV1 (H–N distance) and CV2 (O–S distance). **(A)** Occupancy map showing the frequency of sampled configurations, highlighting a narrow, continuous pathway connecting reactant- and product-like regions. **(B)** Same CV space coloured by simulation time, illustrating the directional progression from larger to shorter CV values along the reaction coordinate. **(C)** CV space coloured by accumulated work, showing that higher work values are associated with configurations in the upper region of the pathway, corresponding to the transition-state region.

### Supporting References

(1) Malojcic, G.; Owen, R. L.; Grimshaw, J. P.; Brozzo, M. S.; Dreher-Teo, H.; Glockshuber, R. A structural and biochemical basis for PAPS-independent sulfuryl transfer by aryl sulfotransferase from uropathogenic Escherichia coli. *Proc Natl Acad Sci U S A* **2008**, *105* (49), 19217-19222. DOI: 10.1073/pnas.0806997105.

(2) Information, N. C. f. B. *NCBI Multiple Sequence Alignment Viewer*. U.S. National Library of Medicine, 2026. https://www.ncbi.nlm.nih.gov/projects/msaviewer/ (accessed April 30, 2026).

(3) Camacho, C.; Coulouris, G.; Avagyan, V.; Ma, N.; Papadopoulos, J.; Bealer, K.; Madden, T. L. BLAST+: architecture and applications. *BMC Bioinformatics* **2009**, *10*, 421. DOI: 10.1186/1471-2105-10-421.

(4) Lua, R. C.; Wilson, S. J.; Konecki, D. M.; Wilkins, A. D.; Venner, E.; Morgan, D. H.; Lichtarge, O. UET: a database of evolutionarily-predicted functional determinants of protein sequences that cluster as functional sites in protein structures. *Nucleic Acids Res* **2016**, *44* (D1), D308-312. DOI: 10.1093/nar/gkv1279.

(5) Krieger, E.; Vriend, G. YASARA View - molecular graphics for all devices - from smartphones to workstations. *Bioinformatics* **2014**, *30* (20), 2981-2982. DOI: 10.1093/bioinformatics/btu426.

(6) Case, D. A.; Aktulga, H. M.; Belfon, K.; Cerutti, D. S.; Cisneros, G. A.; Cruzeiro, V. W. D.; Forouzesh, N.; Giese, T. J.; Gotz, A. W.; Gohlke, H.; et al. AmberTools. *J Chem Inf Model* **2023**, *63* (20), 6183-6191. DOI: 10.1021/acs.jcim.3c01153.

(7) Chovancova, E.; Pavelka, A.; Benes, P.; Strnad, O.; Brezovsky, J.; Kozlikova, B.; Gora, A.; Sustr, V.; Klvana, M.; Medek, P.; et al. CAVER 3.0: a tool for the analysis of transport pathways in dynamic protein structures. *PLoS Comput Biol* **2012**, *8* (10), e1002708. DOI: 10.1371/journal.pcbi.1002708.

(8) Schrodinger, LLC. The PyMOL Molecular Graphics System, Version 1.8. 2015.

(9) Mongan, J.; Case, D. A.; McCammon, J. A. Constant pH molecular dynamics in generalized Born implicit solvent. *J Comput Chem* **2004**, *25* (16), 2038-2048. DOI: 10.1002/jcc.20139.

(10) Schrodinger, LLC. The AxPyMOL Molecular Graphics Plugin for Microsoft PowerPoint, Version 1.8. 2015.

(11) Roe, D. R.; Cheatham, T. E., 3rd. PTRAJ and CPPTRAJ: Software for Processing and Analysis of Molecular Dynamics Trajectory Data. *J Chem Theory Comput* **2013**, *9* (7), 3084-3095. DOI: 10.1021/ct400341p.

(12) Tian, C.; Kasavajhala, K.; Belfon, K. A. A.; Raguette, L.; Huang, H.; Migues, A. N.; Bickel, J.; Wang, Y.; Pincay, J.; Wu, Q.; et al. ff19SB: Amino-Acid-Specific Protein Backbone Parameters Trained against Quantum Mechanics Energy Surfaces in Solution. *J Chem Theory Comput* **2020**, *16* (1), 528-552. DOI: 10.1021/acs.jctc.9b00591.

(13) Michaud-Agrawal, N.; Denning, E. J.; Woolf, T. B.; Beckstein, O. MDAnalysis: a toolkit for the analysis of molecular dynamics simulations. *J Comput Chem* **2011**, *32* (10), 2319-2327. DOI: 10.1002/jcc.21787.

(14) Gaus, M.; Cui, Q.; Elstner, M. DFTB3: Extension of the self-consistent-charge density-functional tight-binding method (SCC-DFTB). *J Chem Theory Comput* **2012**, *7* (4), 931-948. DOI: 10.1021/ct100684s.

(15) Jarzynski, C. Nonequilibrium Equality for Free Energy Differences. *Physical Review Letters* **1997**, *78* (14), 2690-2693. DOI: 10.1103/PhysRevLett.78.2690.
